# From Diminishing Returns to Entrenchment: A Unifying Theory of Epistasis along Adaptive Walks Revealed by Fisher’s Geometric Model

**DOI:** 10.64898/2026.05.07.723117

**Authors:** Arnaud Maillard, Antoine Maillard, Ilan Coeuille, Olivier Tenaillon

**Affiliations:** Institut Cochin, Université Paris Cité, CNRS, INSERM, Paris, France; INRIA, Paris, France; Departement d’Informatique, Ecole Normale Superieure, PSL & CNRS, Paris, France

**Keywords:** Fisher’s geometric model, epistasis, adaptive walk, antagonistic pleiotropy, entrenchment, diminishing returns epistasis

## Abstract

Epistasis makes the fitness effect of a mutation depend on the genetic background in which it occurs, thereby shaping the accessibility and reversibility of evolutionary trajectories. Along adaptive walks, this path-dependent epistasis can take distinct forms: contingency, when a mutation requires prior substitutions to be beneficial; entrenchment, when later substitutions make its reversion increasingly deleterious; and diminishing returns, when successive beneficial mutations reduce one another’s effects. Although these regimes have been documented experimentally, the conditions under which each predominates remain poorly understood. Here we use Fisher’s geometric model to derive a general framework for path-dependent epistasis under stabilizing selection on a multidimensional phenotype. We show that epistasis between substitutions has a simple geometric interpretation: contingency and entrenchment arise when the collateral effects of mutations, orthogonal to the direction of the optimum, are compensated by the preceding or subsequent adaptive path, whereas diminishing returns arise when successive substitutions remain strongly aligned with the same direction of selection. Analytical results and simulations reveal a transition controlled by a single composite parameter combining phenotypic complexity, mutation size, and distance to the optimum. Far from the optimum, adaptive walks are dominated by diminishing returns epistasis. As populations approach the optimum, or as phenotypic complexity increases, antagonistic pleiotropy generates systematic contingency and entrenchment. At mutation-selection-drift equilibrium, these effects become strong, rapidly established, and increase with phenotypic complexity. These results show that contingency and entrenchment do not require specific molecular interactions between residues: they emerge generically from nonspecific epistasis produced by stabilizing selection on pleiotropic traits. Fisher’s geometric model thus unifies diminishing returns, contingency, and entrenchment as distinct regimes of the same underlying geometry of adaptation.

## Introduction

The concept of fitness landscape, introduced by Sewall Wright in 1932 (Wright et al. 1932), provides a powerful framework for understanding the dynamics of adaptive evolution. Formally, a fitness landscape is defined by a set of genotypes, the mutational distances between them, and their associated fitness values. Each genotype is mapped to a scalar, defining a surface over the high-dimensional space of all possible genotypes in which mutational neighbors occupy adjacent positions (De Visser and Krug 2014; Fragata et al. 2019). This geometric representation offers an intuitive picture of evolutionary dynamics. For instance, the distribution of mutational fitness effects at any given genotype is entirely determined by the local curvature of the landscape around that genotype. The evolution of a population can be conceived as a walk through genotype space, driven by natural selection toward regions of higher fitness (Fragata *et al*. 2019).

Wright’s motivation for introducing the fitness landscape was his conviction that, contrary to Fisher’s additive view of genetics, real genotype-fitness maps are complex owing to pervasive genetic interactions, or epistasis. By definition, fitness epistasis occurs when allelic variation at one locus affects the fitness differences conferred by alleles at other loci (Bank 2022). Epistasis is said to be positive if the combination of variants has a higher fitness than expected from their individual effects, and negative otherwise (Domingo *et al*. 2019). Without epistatic interactions, the fitness effect of each mutation would be an intrinsic property of that mutation alone, independent of genetic background, and the landscape would reduce to a trivial additive surface that is smooth, single-peaked, and of limited evolutionary interest. With epistasis, the fitness effect of a mutation is therefore not a fixed quantity, but a function of the genetic background in which it arises. In terms of evolutionary dynamics, epistasis generates multiple fitness peaks (Bank 2022) and constrains the mutational pathways accessible under strong selection (Weinreich *et al*. 2006). The empirical reality of pervasive epistasis has since provided strong justification for the theoretical development of adaptive landscape models. Both experimental data from diverse biological systems (Lunzer *et al*. 2010; Birgy *et al*. 2026; Dixon and Dixon 2004) and comparative genomic data (Breen et al. 2012; Naumenko et al. 2012; Vigué et al. 2022) demonstrate that mutational effects are fundamentally context-dependent. A striking illustration is the observation that derived amino acid states fixed in one species can be severely deleterious, or even lethal, when introduced into a closely related one (Kulathinal et al. 2004).

Beyond shaping the global topology of the fitness landscape, epistasis modifies the mutational spectrum accessible to a population at each successive step along an evolutionary trajectory (Bank 2022; Weinreich et al. 2006). A mutation that is beneficial at the time of its introduction may confer its beneficial effect only in the presence of prior permissive substitutions (for example, a stabilizing mutation) (Ortlund et al. 2007). The fate of a mutation arising in a population may thus be contingent on previous mutations (Bloom et al. 2010; Harms and Thornton 2014). This phenomenon is referred to as evolutionary contingency. Conversely, once a mutation has fixed, it becomes part of the genetic background onto which subsequent modifications are introduced. If the fitness effects of those subsequent substitutions depend on the focal mutation, its reversion becomes increasingly deleterious over time (Bridgham et al. 2009). This phenomenon is called entrenchment. Contingency and entrenchment both reflect positive epistasis between the focal mutation and other substitutions along the trajectory. They may involve not only pairwise interactions but also higher-order epistasis with the full set of accumulated substitutions (Shah et al. 2015). The opposite regime arises under diminishing returns epistasis. A mutation that was strongly beneficial at the time of its fixation may later become dispensable, or even reversible without fitness cost, once the population has further adapted (Chou *et al*. 2011; Khan et al. 2011). This reflects negative epistasis between successive beneficial substitutions. Critically, these phenomena concern epistatic interactions specifically among mutations that fix along a trajectory, which are a strongly non-random subset of all possible mutations. We use *path-dependent epistasis* to refer to all forms of epistasis in which mutational effects depend on prior evolutionary history, including contingency, entrenchment, and temporal shifts in fitness effects.

Although path-dependent epistasis is an inherent property of any fitness landscape with epistatic interactions, its formalization and characterization have been driven primarily by experimental work rather than theoretical developments. Evidence for contingency (Blount et al. 2018), entrenchment, and diminishing returns epistasis (de Visser et al. 2011) has accumulated across biological systems of very different levels of biological complexity. The Long-Term Evolution Experiment (LTEE), in which replicate populations of *Escherichia coli* have been evolving under identical conditions for more than 80,000 generations, provides compelling illustrations at the level of whole-organism fitness. The evolution of aerobic citrate utilization in a single LTEE population, a novel metabolic capacity absent from all other replicates after more than 30,000 generations, was shown to depend on prior permissive substitutions that potentiated the trait, exemplifying evolutionary contingency at the genomic scale (Blount et al. 2008). The same experiment has documented pervasive diminishing returns epistasis among successive beneficial substitutions (Khan et al. 2011), consistent with negative epistasis along adaptive trajectories. The most mechanistically detailed evidence, however, comes from proteins, whose function is tightly coupled to three-dimensional structure, making the context-dependence of amino acid substitutions both expected and experimentally tractable (Starr and Thornton 2016). Comparing substitutions fixed between orthologs maintaining the same function either after experimental evolution or in closely related species provides a first class of evidence. Lunzer *et al*. (2010) introduced each of the 168 amino acid differences separating *Escherichia coli* and *Pseudomonas aeruginosa* isopropylmalate dehydrogenase individually into the heterologous background. Approximately one third severely compromised enzymatic activity, demonstrating that permissive or restrictive substitutions in each lineage had rendered these states strongly context-dependent. Ancestral protein reconstruction has provided perhaps the most direct window into path-dependent epistasis. In the glucocorticoid receptor, the evolution of cortisol specificity required prior permissive substitutions that stabilized specific structural elements, allowing the protein to tolerate subsequent function-switching mutations (Ortlund *et al*. 2007). These permissive substitutions were found to be exceedingly rare among the accessible sequence variants (Harms and Thornton 2014), underscoring the strong contingency of the transition. In one of the most comprehensive quantitative assessment to date, Starr *et al*. (2018) traced the evolutionary history of the Hsp90 ATPase domain over one billion years. They found that more than 75% of historical substitutions were either contingent on prior permissive substitutions, entrenched by subsequent restrictive ones, or both, a result independently confirmed by Park et al. across 700 million years of steroid receptor evolution (Park *et al*. 2022). Finally, at a broader scale, analyses of multiple sequence alignments of homologous proteins using statistical physics-based models such as Direct Coupling Analysis (DCA) (Weigt *et al*. 2009; Cocco *et al*. 2018) have shown that incorporating pairwise epistasis substantially improves the prediction of mutational effects (Figliuzzi *et al*. 2016; Chen *et al*. 2024; Birgy *et al*. 2026) and, crucially, enables the generation of functional sequences (Russ *et al*. 2020) — something approaches neglecting epistasis consistently fail to achieve. In line with ancestral reconstruction experiments, these studies further indicate that, at the proteome-wide level, a substantial fraction of mutations — on the order of 30 to 50% — have effects that are strongly context-dependent (Vigué *et al*. 2022). Taken together, these studies establish that path-dependent epistasis is not a rare curiosity but a pervasive feature of molecular evolution, shaping accessible trajectories over timescales from laboratory experiments to deep evolutionary time.

Despite the compelling evidence reviewed above, several fundamental questions about path-dependent epistasis remain unresolved. A first question concerns prevalence across proteins and evolutionary timescales. Permissive mutations appear to be rare in the sequence neighborhoods of extant proteins (Harms and Thornton 2014), yet their functional consequences can be dramatic. More broadly, negative epistasis between random mutation pairs consistently outnumbers positive epistasis in deep mutational scanning studies (Starr and Thornton 2016; Kemble *et al*. 2019). This raises the question of what fraction of substitutions fixed along adaptive trajectories actually experience the positive epistasis underlying contingency and entrenchment, rather than the negative epistasis underlying diminishing returns among beneficial substitutions. A second question concerns the conditions under which each regime predominates. Evidence for entrenchment comes primarily from proteins near mutation-selection-drift equilibrium, such as Hsp90, which has been evolving under long-term purifying selection. Evidence for diminishing returns epistasis, in contrast, comes primarily from experimental adaptation to novel selective constraints. This contrast suggests the following hypothesis: diminishing returns epistasis may dominate early in adaptation, when populations are far from a fitness optimum and successive beneficial mutations antagonize one another, while entrenchment becomes the dominant regime as populations approach the optimum and neutral substitutions progressively restrict the reversibility of earlier ones. Whether this reflects a general principle or an artifact of the limited and contrasting experimental systems available remains an open question. A third question concerns mechanistic basis. Starr and Thornton (2016) distinguished two broad classes of epistatic interaction. In specific or microscopic epistasis, a mutation modulates the effects of a small number of other mutations through direct physical interactions between residues. For instance, one substitution may be deleterious due to steric hindrance unless the adjacent amino acid is first replaced. In nonspecific epistasis, sometimes called global (Otwinowski *et al*. 2018) or macroscopic epistasis (Good and Desai 2015), mutations are additive with respect to an underlying physical property but interact nonadditively at the level of fitness. This arises from a nonlinear mapping between the physical property and fitness. The canonical example is the sigmoidal relationship between protein stability and fraction folded (Starr and Thornton 2016; Kemble et al. 2019). The evolutionary implications differ sharply. According to some authors (Starr and Thornton 2016), specific epistasis tends toward positive interactions, rare permissive mutations, and strong historical contingency. Non-specific epistasis tends toward negative interactions, broadly distributed permissive effects, and weaker contingency. A key open question is therefore whether path-dependent epistasis in long-term evolution reflects primarily specific, structurally idiosyncratic interactions, or emerges generically from nonlinear genotype-phenotype mappings. This distinction has direct consequences for the generality and predictability of evolutionary entrenchment and contingency.

Addressing these questions requires frameworks that can move beyond individual case studies toward general, quantitative predictions. Several complementary approaches have been pursued. Empirical fitness landscapes provide direct access to epistatic structure, and their scope has expanded with high-throughput techniques (Kemble et al. 2019), but the combinatorial explosion of genotype space limits them to a handful of mutations and their generalizability remains unknown. Ancestral sequence reconstruction offers a temporally resolved view of path-dependent epistasis, but is restricted to single realized paths, most often involving functional transitions, and becomes intractable with large numbers of substitutions. Statistical models inferred from sequence alignments, such as DCA, capture the epistatic architecture of protein families at large scale, but offer limited mechanistic interpretability (Schmelkin et al. 2025). Finally, mechanistic in silico simulations, including lattice models or stability-based approaches such as FoldX (Shah et al. 2015), provide a direct link between sequence, structure, and fitness, but are restricted to purifying selection on thermodynamic stability, leave adaptive trajectories largely unexplored, and rely on assumptions of uncertain biological realism and generalizability.

Theoretical models of adaptive landscapes offer a complementary approach, abstracting the biological system to gain in generality and intelligibility (Servedio et al. 2014). Among these, Fisher’s geometric model (FGM) is particularly well-suited to investigate path-dependent epistasis along adaptive trajectories. Originally proposed in 1930 (Fisher 1930), FGM is a phenomeno logical model in which individuals are characterized by **n** continuous phenotypic traits under stabilizing selection toward a single Gaussian fitness optimum, each trait corresponding to an axis in an n-dimensional Euclidean space (Tenaillon 2014). Despite its simplicity, FGM approximates a broad class of systems biology models and has been shown to emerge from relatively general first principles describing the underlying metabolic networks and developmental processes of organisms (Martin 2014). It naturally captures the simultaneous optimization of multiple traits: for a protein-coding gene, catalytic activity, RNA stability, translation efficiency, folding speed, interaction with molecular partners, and avoidance of misfolding and aggregation all contribute jointly to fitness (DePristo *et al*. 2005). With just three parameters, phenotypic complexity ***n***, distance to the optimum ***d***, and mutational size ***r***, FGM generates quantitative predictions on the distribution of fitness effects, the rate of adaptation, and the distribution of effect sizes (Orr 2000, 2006; Martin and Lenormand 2006).

Epistasis has been studied extensively in this model (Martin *et al*. 2007; Gros *et al*. 2009; Blanquart *et al*. 2014; Hwang *et al*. 2017). Epistasis arises in FGM as an emergent property of the curvature of the fitness surface: the same phenotypic displacement produces a fitness effect that depends on the starting position. While the distribution of epistasis between two random mutations is centered around zero, prior work has established that epistasis among beneficial mutations from the same origin is negative on average, because such mutations tend to point toward the optimum and thus follow the concave curvature of the fitness surface (Martin *et al*. 2007). This might at first glance suggest that FGM generates only diminishing returns epistasis along adaptive trajectories. The critical question, however, concerns epistasis between two successive mutations along a walk. The first substitution shifts the phenotype and thereby alters the selective context experienced by the second, making this a substantially harder problem. Analytical results exist only in the limiting case of mutations arising very far from the optimum (Blanquart *et al*. 2014), and no general solution is available. The problem is further compounded when considering path-dependent epistasis in the full sense, where the fitness effect of reverting or adding any mutation depends on the entire accumulated genetic context, potentially involving higher-order interactions among all prior substitutions. FGM nonetheless remains tractable in this setting, because the fitness effect of adding or removing a mutation depends only on the initial and final phenotypic positions, not on the intermediate steps of the walk. Here we exploit this tractability to investigate systematically whether contingency, entrenchment or diminishing return can emerge as generic properties of adaptive walks in FGM, and under what conditions each regime predominates.

### Materials and methods

### Fisher’s Geometric Model

Individual fitness depends on an organism’s phenotypic state across *n* quantitative traits under stabilizing selection. A phenotype is represented by a vector **z** = (*z*^1^, …, *z*^*n*^) ∈ ℝ^*n*^, where superscripts index phenotypic components. We denote by ⟨·, ·⟩the canonical inner product on ℝ^*n*^ and by∥ · ∥ the associated Euclidean norm.

Each mutation is represented by a vector ***δ*** ∈ ℝ^*n*^. An evolutionary trajectory is a sequence (**z**_*i*_)_*i*≥ 0_ starting from an initial phenotype **z**_0_, with

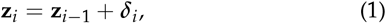

where subscripts index substitution steps along the adaptive walk. The distance to the optimum at step *i* is *d*_*i*_:= ∥**z**_*i*_∥.

We adopt three standard assumptions of Fisher’s geometric model (FGM) (Tenaillon 2014). First, each phenotypic component is subject to stabilizing selection, and fitness is a smooth, single-peaked function of phenotypic position. Without loss of generality, the phenotypic optimum is placed at the origin of ℝ^*n*^, with axes taken as orthogonal. Each component *z*^*k*^ thus measures the deviation of the organism from the optimal value at trait *k*.

Second, fitness takes the Gaussian form

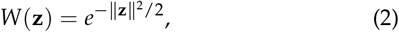

so that log-fitness is

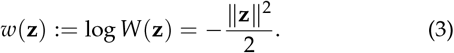

Third, mutations are assumed to affect all phenotypic traits simultaneously (universal pleiotropy). Each mutational vector ***δ*** is drawn independently from an isotropic multivariate Gaussian distribution with mean zero and variance *σ*^2^ per axis:

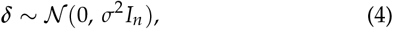

where *I*_*n*_ is the *n* × *n* identity matrix.

For an isotropic Gaussian mutation, ∥***δ***∥ follows a chi distribution with *n* degrees of freedom, with expectation

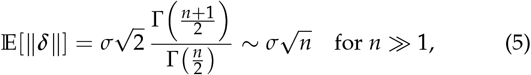

and variance

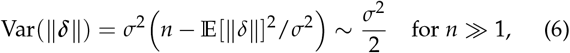

so that for large *n*, ∥***δ***∥ concentrates around 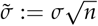with fluctuations of order *σ*. For notational clarity, we write *r*:= ***δ*** for the mutational size, and use either *r* or ***δ*** throughout, depending on which makes the geometric intuition or connection with prior literature more transparent.

The log-fitness effect of a mutation ***δ*** arising from phenotypic position **z** is

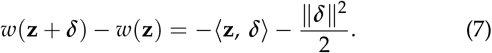

The first term, −⟨**z, *δ***⟩, captures the directional component: a mutation moving toward the optimum (i.e., with ⟨**z, *δ*** ⟩ *<* 0) increases log-fitness. The second term, −∥***δ***∥^2^/2, is a curvature penalty that is always negative and grows with mutational size. A mutation is beneficial if and only if equation (7) is positive, which requires ⟨**z, *δ***⟩ *<*− ∥***δ***∥^2^/2, or equivalently cos *θ <* −*r*/(2*d*), where *θ* = ∠(***δ*, z**), *r* = ∥***δ***∥ and *d* = ∥**z**∥ (Figure 1).

**Figure 1.**
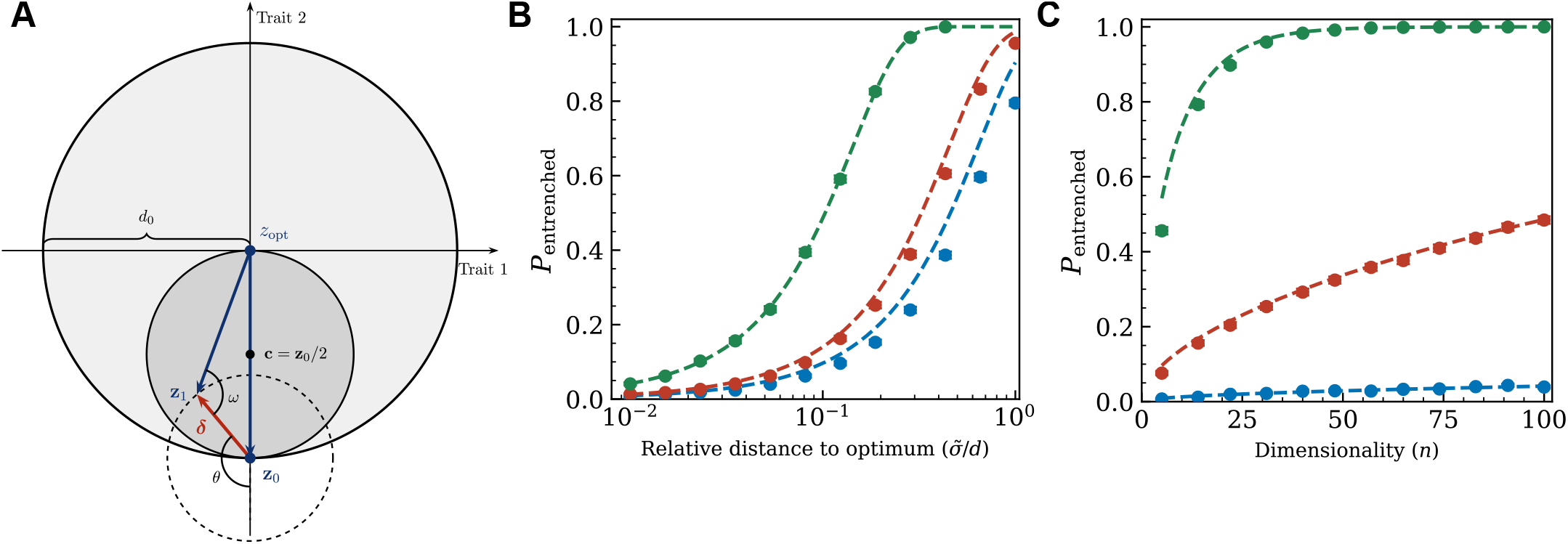
Geometry of entrenchment in Fisher’s geometric model and probability that a beneficial substitution is entrenched at the optimum. **(a)** A single mutational step ***δ*** arising from phenotype **z**_0_, at distance *d*_0_ = ∥**z**_0_∥from the optimum *z*_opt_. The outer circle (radius *d*_0_, centered at **z**_opt_) delimits the beneficial region: a mutation is beneficial if and only if **z**_0_ + ***δ*** lies within this circle, equivalently if cos *θ <*− ∥***δ***∥ /(2*d*_0_), where *θ* = ∠(***δ*, z**_0_). The inner gray sphere (radius *d*_0_/2, centered at **z**_0_/2) defines the boundary between two epistatic regimes. If **z**_0_ + ***δ*** lands inside the inner sphere, the substitution falls in the diminishing returns regime: its reversion at the optimum would cost less fitness than it originally gained. If **z**_0_ + ***δ*** lands outside the inner sphere but within the outer circle — as illustrated — the substitution is entrenched: its reversion at the optimum becomes more costly than its original benefit. The angle *π* − *ω* denotes the angle between ***δ*** and the new selection axis −**z**_1_:= −(**z**_0_ + ***δ***); entrenchment occurs when *ω < π*/2, i.e. when ***δ*** is anti-aligned with the subsequent direction of selection. **(b)** Probability of entrenchment *P*_entrenched_ as a function of the typical mutational norm relative to the distance to the optimum, 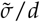, where 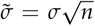 is the expected mutational norm and *d* is the distance to the optimum (log scale on the *x*-axis, linear on the *y*-axis). Dashed curves show the theoretical prediction from equation 18, for *n* = 5 (blue), *n* = 10 (red) and *n* = 100 (green). Symbols show estimates from 10^4^ simulated beneficial mutations drawn from 𝒩(0, *σ*^2^ *I*_*n*_) with Wilson 95% confidence intervals; when error bars are not visible, they are smaller than the symbol. **(c)** Probability of entrenchment *P*_entrenched_ shown as a function of the number of phenotypic dimensions *n* (linear scale on both axes), for 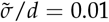 (blue), 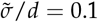 (red) and 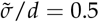 (green). Dashed curves and symbols as in **(b)**.

### Simulation of adaptive walks

To investigate path-dependent epistasis, we simulated adaptive walks under the FGM described above. We assumed that effective population size *N*_*e*_ is large and genomic mutation rate *µ* is small, such that *N*_*e*_*µ* ≪1. Under this strong-selection weak-mutation (SSWM) regime (Gillespie 1991), the population is nearly always monomorphic, and adaptation proceeds through the sequential fixation of new mutations arising in a genetically homogeneous background.

We considered three simulation scenarios. In the first, populations adapt toward the optimum from a distant initial phenotype **z**_0_. Because deleterious mutations have negligible fixation probability far from the optimum, they are ignored; each substitution is drawn uniformly at random from the pool of beneficial mutations. This scenario corresponds to the analytically tractable framework developed in this paper. In the second scenario, only beneficial mutations are considered, but each is accepted with probability

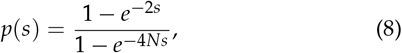

where *s* is the selection coefficient, allowing the full dependence of fixation probability on selection intensity to be accounted for. In the third scenario, we simulated populations at mutation-selection-drift equilibrium (MSDE). Both deleterious and beneficial mutations can fix with probability *p*(*s*), and the population reaches a stationary distribution of phenotypic states. Under the standard FGM with Gaussian fitness, the stationary distribution of log-fitness follows a Gamma distribution, reflecting the drift load imposed by the balance between deleterious and compensatory substitutions (Sella 2009; Tenaillon *et al*. 2007).

Mutational vectors were simulated by drawing random phenotypic displacements from 𝒩 (0, *σ*^2^ *I*_*n*_) and retaining each vector according to the relevant acceptance criterion (rejection sampling). However, rejection sampling becomes computationally prohibitive near the optimum or at large phenotypic dimensionality *n*: the probability of drawing a beneficial mutation decreases super-exponentially as *n* increases or as *d*→ 0. We therefore implemented a Metropolis-Hastings MCMC algorithm to sample efficiently from the exact target distribution, defined as the distribution of mutational vectors ***δ*** conditional on beneficiality and, where applicable, on fixation probability *p*(*s*(***δ***)). This approach remains tractable regardless of dimensionality or proximity to the optimum. Code is available at [GitHub link]. Figure S1 demonstrates the equivalence of mutational vector distributions produced by MCMC and rejection sampling.

### Scaling of mutational effects with phenotypic dimensionality

Because mutations affect *n* phenotypic dimensions simultaneously, their expected norm grows with dimensionality: 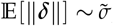, where 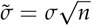 is the typical mutational norm. To compare mutational properties across values of *n* while holding the mean fitness effect of random mutations, equal to 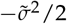 (Tenaillon 2014), approximately constant, we fix 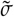 and scale the mutational standard deviation as

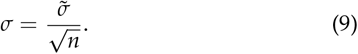

All simulations use this scaling; our analytical results are otherwise general with respect to the relationship between *σ* and *n*.

A key geometric consequence of this scaling concerns the angular distribution of mutational vectors. As *n* increases, the distribution of mutational norms concentrates around 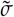. Simultaneously, mutational vectors become increasingly oriented orthogonally to the axis connecting the current phenotype **z** to the optimum. This equatorial concentration is a classical property of high-dimensional geometry, well documented in the context of Fisher’s geometric model (Hartl and Taubes 1996; Poon and Otto 2000): in high dimensions, random vectors are almost surely nearly orthogonal to any fixed direction. When a mutation affects *n* traits independently, its projection onto any fixed direction — including that of the optimum — becomes indeed negligible relative to its total norm as *n* increases. Formally, the angle *θ* = ∠(***δ*, z**) (resp. cos *θ*) concentrates around *π*/2 (resp. 0) as a normal distribution with standard deviation of order 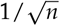 (see Supplementary Materials). A direct consequence is that mutations pointing directly toward the optimum become extremely rare at high complexity, since the beneficial condition cos *θ <* −*r*/(2*d*) requires *θ* to deviate substantially from *π*/2 in the direction of the optimum.

## Results

### Generalized epistasis coefficient

Epistasis between two mutations is defined as the deviation of the log-fitness effect of the double mutant from the sum of the log-fitness effects of the single mutants:

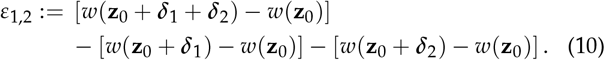

As emphasized by Shah *et al*. (2015), this can equivalently be written as the difference between the fitness effect of mutation ***δ***_2_ in the background carrying mutation ***δ***_1_ and its effect in the naive background:

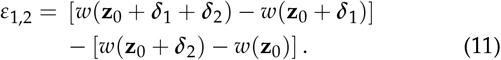

This interpretation of epistasis in terms of substitution order suggests a natural generalization to longer evolutionary trajectories, initially proposed by Shah *et al*. (2015). Consider a trajectory fixing substitutions ***δ***_1_, ***δ***_2_, … sequentially, with 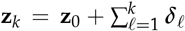. For any pair of substitutions *i* and *j*, one can ask how much larger the fitness effect of substitution ***δ***_*i*_ would have been had it occurred at position *j* along the trajectory rather than at position *i*. When *i > j*, this compares the fitness effect of ***δ***_*i*_ in an earlier background **z**_*j* −1_ to its actual effect at fixation in **z**_*i* −1_. When *i < j*, this is equivalently interpreted as asking whether reverting ***δ***_*i*_ has become more costly in the later background **z**_*j*_ than it was beneficial at fixation, reflecting epistasis with the intervening substitutions ***δ***_*i*+1_, …, ***δ***_*j*_. The generalized epistasis coefficient between substitution *i* and position *j* is defined as

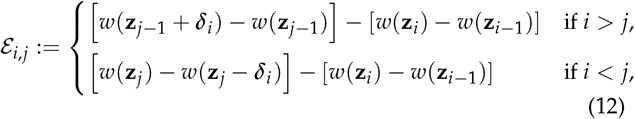

where *w*(**z**_*k*_) denotes the log-fitness of the genotype carrying substitutions ***δ***_1_ through ***δ***_*k*_. Note that ℰ_*i,i*+1_ reduces to the standard pairwise epistasis between two consecutive substitutions.

This generalized measure provides a natural framework for defining contingency and entrenchment. Substitution ***δ***_*i*_ is said to be *contingent* on preceding substitutions if, for *j < i*, ℰ_*i,j*_ *<* 0: the fitness effect of ***δ***_*i*_ would have been smaller, or even negative, had it arisen earlier in the trajectory. Conversely, substitution ***δ***_*i*_ is said to be *entrenched* by subsequent substitutions if, for *j > i*, ℰ_*i,j*_ *>* 0: reverting ***δ***_*i*_ becomes increasingly deleterious as later substitutions accumulate.

Epistasis takes a particularly simple form in FGM, with a strong geometric interpretation. Consider two mutations ***δ***_1_ and ***δ***_2_ fixing sequentially in an individual of phenotype **z**_0_: ***δ***_1_ fixes first, bringing the phenotype to **z**_0_ + ***δ***_1_, and ***δ***_2_ then fixes in this new background. Assuming phenotypic additivity of mutations, a direct calculation using *w*(**z**) = −∥**z**∥^2^/2 yields

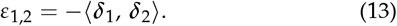

Epistasis is thus negative when the two mutations point in the same direction, zero when they are orthogonal, and positive when the angle between them exceeds *π*/2.

Consider now an adaptive walk in FGM. Starting from phenotype **z**_0_, a sequence of mutations fixes successively, so that 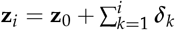. Extending the framework above, we show in Supplementary Materials that the generalized epistasis coefficient takes an analogous simple form. For *i* ≥ *j*:

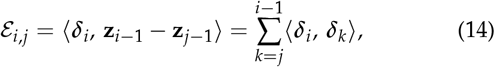

and for *i < j*:

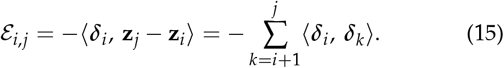

The interpretation is analogous to that of pairwise epistasis. For *i*≥ *j*, ℰ_*i,j*_ measures the alignment between ***δ***_*i*_ and the segment **z**_*i* −1_− **z**_*j* −1_, which represents the phenotypic displacement accumulated by substitutions ***δ***_*j*_, …, ***δ***_*i* −1_ prior to the fixation of ***δ***_*i*_. When this earlier path is aligned with ***δ***_*i*_, ℰ_*i,j*_ is positive: ***δ***_*i*_ would have been more beneficial at step *j* than when it actually occurred at step *i*, a case of diminishing returns. Substitution ***δ***_*i*_ is contingent on substitutions ***δ***_*j*_, …, ***δ***_*i* −1_ when ℰ_*i,j*_ *<* 0: it would have been less beneficial, or deleterious, before those substitutions occurred.

For *i < j*, ℰ_*i,j*_ measures the alignment between ***δ***_*i*_ and the sub-sequent mutational path **z**_*j*_− **z**_*i*_. Substitution ***δ***_*i*_ is entrenched (ℰ_*i,j*_ *>* 0) when subsequent substitutions tend to point in the opposite direction to ***δ***_*i*_: they collectively compensate its phenotypic effect, making reversion increasingly costly. Conversely, ℰ _*i,j*_ *<* 0 when ***δ***_*i*_ is aligned with the subsequent path, a case of diminishing returns epistasis in which later substitutions reduce the fitness contribution of ***δ***_*i*_.

### Entrenchment of early substitutions in an adaptive walk toward the optimum

#### Geometric interpretation

We now consider the limiting case in which the adaptive walk reaches the phenotypic optimum, which provides a clear geometric intuition for entrenchment. Starting from phenotype **z**_0_, suppose that mutation ***δ***_1_ fixes first, bringing the phenotype to **z**_1_ = **z**_0_ + ***δ***_1_. The adaptive walk then continues with subsequent substitutions that bring the phenotype arbitrarily close to the optimum, so that **z**_∞_ = **0**.

Although it is clear that reverting ***δ***_1_ once at the optimum incurs a fitness cost, it is not a priori obvious whether this cost is larger or smaller than the benefit of ***δ***_1_ when it initially fixed at **z**_0_. We show in Supplementary Materials that ℰ_1,∞_ takes a simple form in this limit. Denoting **c** = **z**_0_/2, the entrenchment of ***δ***_1_ depends only on the distance between **c** and **z**_1_ = **z**_0_ + ***δ***_1_:

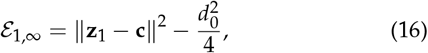

where *d*_0_ = ∥**z**_0_∥ is the initial distance to the optimum. For a given 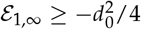, **z**_1_ lies on a sphere of center **c** and radius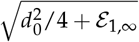.

As shown in Figure 1, the sphere of center **c** = **z**_0_/2 and radius *d*_0_/2 — which passes through both the origin and **z**_0_ — defines a natural boundary between two regimes. If **z**_1_ lands inside this sphere, the mutation falls in a regime of diminishing returns epistasis: reverting ***δ***_1_ at the optimum is less costly than its initial benefit at **z**_0_. If **z**_1_ lands outside this sphere, ***δ***_1_ is entrenched: reverting it at the optimum is more costly than its initial benefit.

The boundary case ℰ_1,∞_ = 0, i.e. **z**_1_ lying exactly on this sphere, corresponds to the case where **z**_1_ is orthogonal to ***δ***_1_. Geometrically, this means that the new selection axis defined by −**z**_1_ is orthogonal to the mutational vector ***δ***_1_. More generally, diminishing returns occurs when the angle *ω* between ***δ***_1_ and **z**_1_ exceeds *π*/2, and entrenchment when this angle is less than *π*/2. Intuitively, a mutation landing outside the central sphere is entrenched because it is anti-aligned with the subsequent selection axis −**z**_1_: the adaptive walk that follows, by moving toward the optimum along this new axis, tends to compensate and correct the phenotypic effect of ***δ***_1_.

#### Probability of entrenchment

In the limiting case where the adaptive walk reaches the optimum, the probability that a beneficial mutation is entrenched can be computed explicitly. Let *θ* denote the angle between ***δ***_1_ and **z**_0_. A mutation of size ***r*** is beneficial if and only if cos *θ <* −*r*/2*d*_0_, and it is entrenched if and only if cos *θ <* −*r*/*d*_0_. Considering fixed ***r*** for simplicity, a classical result of FGM first established by Fisher (Fisher 1930) and recovered by subsequent authors (Hartl and Taubes 1996; Ram and Hadany 2015) shows that for small mutation size and large *n*, the probability of being beneficial depends on a single dimensionless parameter 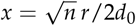:

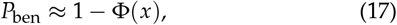

where Φ denotes the cumulative distribution function of the standard normal distribution. In the case of Gaussian mutations considered in this article, the norm ***r*** is itself a random variable, but it concentrates around 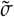 for large *n*. The parameter *x* is thus defined as 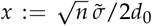 (Tenaillon 2014), and equation (17) still holds.

Intuitively, with a mutation of fixed size ***r***, when the current phenotype is very far from the optimum (small *r*/*d*_0_, i.e. a severely maladapted phenotype), the fraction of beneficial mutations approaches 50%. As phenotypic complexity *n* increases, however, this fraction decreases. Biologically, mutations are more likely to disrupt a complex system than a simple one: with more traits under selection, a random phenotypic displacement is more likely to worsen at least one of them. The dimensionless parameter *x* captures a competition between two geometric effects. The half relative distance to the optimum *y*:= *r*/(2*d*_0_), which appears naturally in the beneficiality condition cos *θ <* −*y* derived from equation (7), controls the curvature of the beneficial region: when *y* is small, the organism is so far from the optimum that the fitness landscape experienced by the mutation appears nearly flat, and almost any displacement is as likely to be beneficial as deleterious. The dimensionality *n*, by contrast, concentrates mutational vectors away from the direction of the optimum: cos *θ* approximately follows a normal distribution with standard deviation 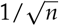 (see Supple-mentary Materials). The beneficial threshold cos *θ <* −*y* thus lies 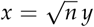 standard deviations below zero, so that *x* jointly captures the effects of relative mutation size and phenotypic complexity.

Returning to the limiting case of an adaptive walk reaching the optimum, the entrenchment condition cos *θ <* −*r*/*d*_0_ is geometrically equivalent to the beneficial condition in a sphere of half the radius, that is, with *x* replaced by 2*x*. The probability of being entrenched given that the mutation is beneficial is therefore:

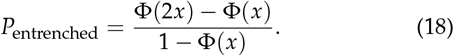

As Figure S2 shows, this probability tends to 0 for small *x*, and increases rapidly toward 1 as *x* increases, as the approximation 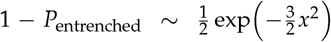 for large *x* indicates. Simulations confirm that equation (18) is accurate for Gaussian mutations and *n* ≥ 5 (Figure S2). Critically, *P*_entrenched_ depends on *n, σ* and *d* only through the composite parameter 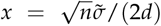: any combination of these three quantities yielding the same value of *x* produces the same probability of entrenchment. Figure 1 illustrates the probability of entrenchment as a function of *n* for several distances to the optimum 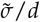. The mutational standard deviation is scaled with *n* such that 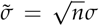 is fixed to assure that the mean fitness effect is constant across dimensionality (see Scaling, Methods). Under this scaling,the probability of entrenchment increases with 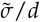 and, in all cases, rises with the number of traits under stabilizing selection. Note the analytical results are general and do not depend on any particular scaling.

We derive in Appendix the expected generalized epistasis 𝔼 [ℰ_1,∞_]:

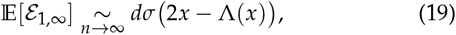

where Λ(*x*) = *ϕ*(*x*)/(1 − Φ(*x*)) is the inverse Mills ratio and *ϕ* denotes the standard normal density. The sign of 𝔼 [ℰ_1,∞_] is controlled by 2*x* − Λ(*x*), and the amplitude at fixed *x* is 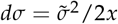. Note that *x* and 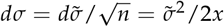 are not independent: *x* combines phenotypic complexity, distance to the optimum, and mutational norm, so that varying *x* necessarily changes the amplitude. The variance satisfies:

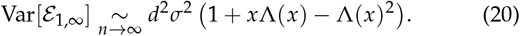

For comparison, the mean log-fitness gain of the first substitution equals

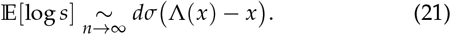

Two biologically distinct regimes emerge (Figure S3-b). Far from the optimum 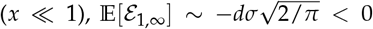: the first substitution undergoes diminishing returns rather than entrenchment. The absolute magnitude depends on how *x* ≪1 is reached: it is large when *d* is large at fixed 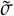 and *n*, and vanishingly small when 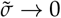 at fixed *d* and *n*. In all cases and most importantly, Figure S3b shows that 𝔼 [ℰ_1,∞_]/ 𝔼 [log *s*] →−1: the expected epistatic cost of reverting the substitution at the optimum approximately cancels its original selective benefit.

Close to the optimum or at high phenotypic complexity 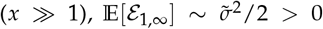. In contrast to the previous regime, this limit is robust: the amplitude depends only on 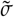, regardless of how *x* ≫ 1 is reached. The coefficient of variation satisfies *CV* ~ 1/*x*^2^ ≪ 1: entrenchment is not only consistently positive but increasingly deterministic. Figure S3b shows that 𝔼 [ℰ_1,∞_]/ 𝔼 [log *s*] diverges as *x*^2^: beneficial mutations concentrate near the boundary of the beneficial sphere, contributing little to immediate fitness while becoming increasingly costly to revert once the population has reached the optimum.

The transition between the two regimes occurs at *x*^∗^ ≈ 0.612, defined by Λ(*x*^∗^) = 2*x*^∗^, where 𝔼 [ℰ_1,∞_] changes sign.

### Epistasis between successive beneficial mutations

Unlike the limiting case ℰ_1,∞_, where the entrenchment of a single substitution is evaluated once the population has reached the optimum, computing path-dependent epistasis along an adaptive walk requires tracking how each substitution modifies both the distance to the optimum and the direction of selection. Entrenchment and contingency of a substitution ***δ***_*i*_ depend respectively on ∑_*k>i*_ ⟨***δ***_*i*_, ***δ***_*k*_ ⟩and ∑_*k<i*_ ⟨***δ***_*i*_, ***δ***_*k*_⟩, that is, on the inner products between ***δ***_*i*_ and the substitutions that follow or precede it along the walk. This sequential dependence makes the problem analytically challenging: the distribution from which each mutation is drawn depends on the entire prior history of the walk.

Analytical results are nonetheless accessible in the simplest case: that of two consecutive substitutions, characterized by ℰ_*i,i*+1_. Two preliminary remarks are in order. First, the definition of generalized epistasis implies directly that ℰ _*i,i*+1_ = −ℰ_*i*+1,*i*_; we therefore consider only ℰ_*i,i*+1_ in what follows. Second, existing results in the literature are restricted to the regime 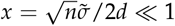, that is, to populations very far from the optimum with sufficiently low phenotypic complexity (Blanquart *et al*. 2014). In this limiting regime, successive substitutions modify the *distance* and *direction* to the optimum only weakly, so that consecutive mutations are drawn from approximately the same distribution. Epistasis between two consecutive substitutions then reduces to epistasis between two beneficial mutations arising in the same genetic background. Since fitness isoclines are nearly planar in this regime, a mutation is beneficial if and only if it has a component pointing toward the optimum. Under these two conditions, Blanquart *et al*. (2014) showed that mean epistasis between two such mutations is negative, reflecting a regime of diminishing returns epistasis. We extend here the calculation of epistasis between two consecutive beneficial substitutions to intermediate and near-optimum regimes.

We derive and present in Supplementary Materials an exact formula for the expected epistasis between two consecutive beneficial mutations, which is complex but numerically tractable as a function of *σ, n*, and *d*. Figure 2-a shows the value of this integral (solid red curve) for *n* = 10, 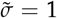 fixed, and varying distance to the optimum parameterized by 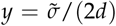. For a mutation arising very far from the optimum, epistasis is on average negative; it then increases and becomes positive, reaching a maximum for *y* between 0.1 and 1. This prediction matches closely the empirical mean obtained from 1 000 simulations of two consecutive beneficial mutations. Simulations further show that the variance is large for small *y* but decreases as *y* increases, reminiscent of what was observed for ℰ_1,∞_.

**Figure 2.**
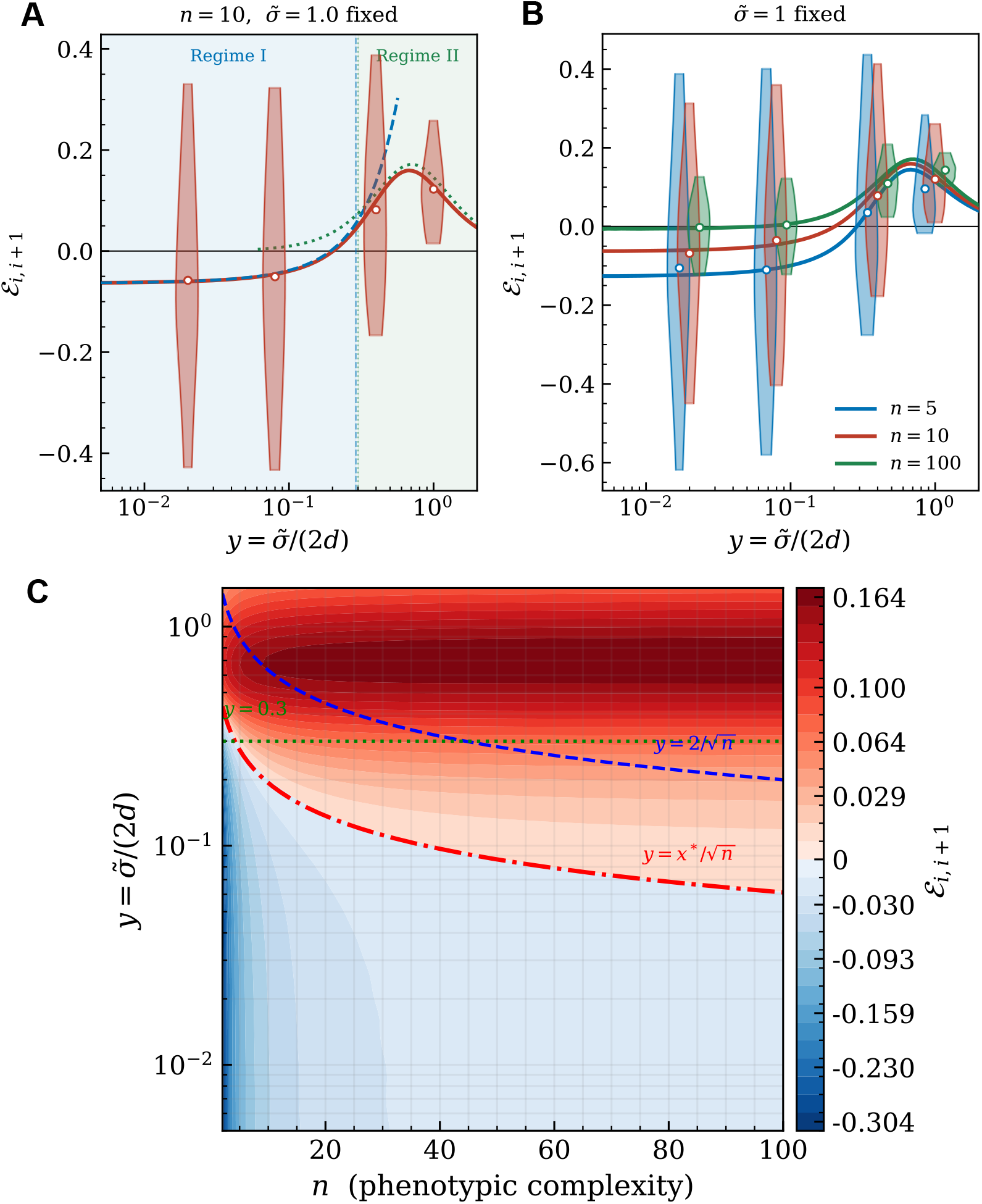
Pairwise epistasisℰ_*i,i*+1_ between two consecutive beneficial mutations. **(a)** ℰ_*i,i*+1_ as a function of 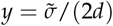, for *n* = 10 and 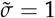 fixed, with 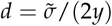 varying. The solid red curve shows the exact numerical prediction. The dashed blue curve shows the Regime I asymptotic 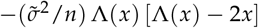 with 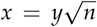, valid for *x* ≤ 1, i.e. 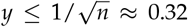 (blue shaded region). The dotted green curve shows the Regime II asymptotic 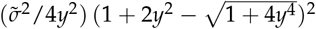, valid for *y* ≥ 0.3 (green shaded region). Violin plots (10th to 90th percentile) show the empirical distribution of ℰ_*i,i*+1_ from 1000 simulated pairs of consecutive beneficial mutations at each value of *y*, drawn without weighting by fixation probability; medians are shown as white dots. In Regime I, ℰ_*i,i*+1_ *<* 0 (diminishing returns); the sign reverses near 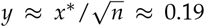 (Recall that *x*\*≈ 0.612). In Regime II, epistasis is systematically positive (entrenchment) and the Regime II asymptotic provides a good approximation for *y* ≥ 0.3. **(b)** Same quantity for *n* = 5, *n* = 10, and *n* = 100 (blue, red, green), with 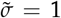 fixed throughout. Solid curves show the exact numerical integral; violin plots show 1000 simulations per (*n, y*) combination (10th to 90th percentile). At fixed 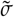 and small *y* (Regime I), the absolute magnitude of epistasis decreases with *n*, consistent with the 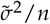 scaling of equation 22. In Regime II (*y* ≥ 0.3), curves for different *n* converge, confirming that the Regime II prediction is independent of dimensionality. The maximum of ℰ_*i,i*+1_ occurs near *y* ≈ 0.7 in all cases. **(c)** Expected pairwise epistasis ℰ_*i,i*+1_ computed from the exact numerical integral across the (*n, y*) plane, with 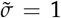 fixed. Color encodes the epistasis value on a signed logarithmic scale truncated at the 98th percentile (red: positive, entrenchment; blue: negative, diminishing returns). The red dash-dotted curve shows the Regime I sign-change boundary 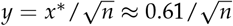, which follows the empirical zero contour precisely. The dashed blue curve shows the outer Regime I boundary 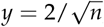, beyond which the Regime I asymptotic becomes inaccurate at moderate *n*. The dotted green horizontal line at *y* = 0.3 marks the lower boundary of Regime II. The epistasis reaches a maximum amplitude near *y* ≈ 0.7, consistent with the analytical prediction 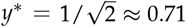 from equation 23.

To better characterize the dynamics of pairwise epistasis, we consider two distinct asymptotic regimes. The first corresponds to large *n* and small *y* of order 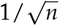, so that 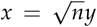 is of order unity — we refer to this as the *far-from-optimum* regime (Regime I). At fixed 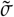, this corresponds to a distance to the optimum of order 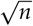. The second regime corresponds to *y* = *O*(1), roughly *y* ∈ [0.3, 1.5] — the *near-optimum* regime (Regime II).

#### Regime I: far from the optimum

This regime is characterized by *x* = *O*(1) and corresponds to a relative mutational size 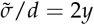 of order 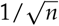. The population is far from the optimum in absolute terms, but the curvature of the fitness landscape influences the distribution of mutational effects in a non-negligible way. In this regime, each substitution appreciably modifies both the distance to the optimum and the direction of selection, so that approximations valid for *x* ≪ 1 no longer apply. We derive analytically that the expected epistasis between two consecutive beneficial mutations takes the form:

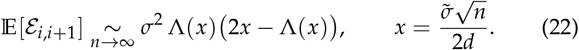

For *x* ≪ 1, this expression reduces to −2*σ*^2^/*π*, a form close to the result of Blanquart *et al*. (2014); the residual difference stems from the fact that we consider here two consecutive beneficial mutations without weighting by fixation probability 2*s*. By symmetry, once ***δ***_*i*_ is fixed, mutations ***δ***_*i*+1_ are uniformly distributed around the new selection axis **z**_*i*_. The geometry is therefore identical to that of ℰ_1,∞_, and the expectation of ℰ_*i,i*+1_ has the same functional form as 𝔼 [ℰ_1,∞_]. One thus recovers the same critical threshold *x*\* ≈ 0.612 separating a regime of negative epistasis (diminishing returns) from a regime of positive epistasis (entrenchment).

For *n* = 10, this approximation is excellent for 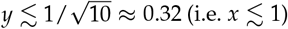, but overestimates the mean epistasis for larger *y*. The sign change predicted by equation (22) occurs at 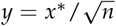, and is perfectly recovered in Figure 2-a. More generally, Figure 2-c shows that the sign change of mean epistasis, computed from the exact integral across the full (*n, y*) plane, follows the boundary 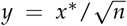 precisely. Note finally that equation (22) involves 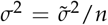: Figure 2b confirms that at fixed 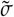 and small *y*, the absolute magnitude of epistasis decreases with *n*, consistent with this 1/*n* scaling. This dependence on *n* disappears in Regime II.

The underlying intuition for the sign change parallels that of the limiting case ℰ_1,∞_. By symmetry, mutations ***δ***_*i*+1_ are distributed around the new selection axis **z**_*i*_, so that the sign of ℰ_*i,i*+1_ is determined by the alignment between ***δ***_*i*_ and **z**_*i*_. For small *x*, beneficial mutations concentrate in the inner sphere (Figure 1): they are on average aligned with **z**_*i*_, generating diminishing returns. For larger *x*, beneficial mutations subtend an angle *θ* closer to *π*/2 with **z**_*i*−1_ and concentrate in the narrow region between the inner and outer spheres, so that ***δ***_*i*_ tends to be anti-aligned with **z**_*i*_ and will on average be compensated by the next substitution, generating entrenchment.

#### Regime II: close to the optimum

Along an adaptive walk, 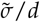 increases as the population approaches the optimum. For a given complexity *n, x* grows accordingly, and the assumption 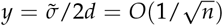 ceases to hold. We therefore consider a second regime defined by *y* = *O*(1), corresponding roughly to 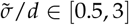. In this regime, beneficial mutations are rare and drawn from the extreme tail of the distribution of mutational effects: their properties are governed by the tail rather than the bulk of the mutational distribution. The expected epistasis between two consecutive beneficial mutations takes the form (see Supplementary Materials):

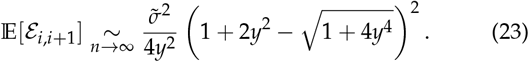

This expression is always positive: in contrast to Regime I, epistasis between consecutive beneficial mutations is systematically positive throughout Regime II, regardless of *n*. The magnitude is of order 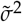, meaning the entrenchment is strong and independent of dimensionality. Figures 2a and 2b confirm that equation (23) provides an excellent approximation for *y* ≳ 0.3 at *n* = 10, and that the curves for different values of *n* converge in this regime, consistently with the *n*-independence of the prediction. Simulations further suggest that variance decreases in this regime.

The function 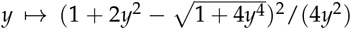 reaches a unique maximum of 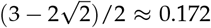 at 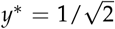, con-sistent with the maximum observed in Figures 2a and 2b. Note that the two regimes are consistent in their common limit (Supplementary Materials).

### Path-dependent epistasis along long adaptive walks

Deriving exact analytical expressions for entrenchment and contingency along longer adaptive walks remains an open problem. We therefore proceed by simulation. The symmetry argument developed above suggests that the sign of mean generalized epistasis over a long walk should be consistent with that obtained for ℰ_*i,i*+1_ and ℰ_1,∞_: the vectors **z**_*j*_ − **z**_*i*_ and **z**_*i*−1_ − **z**_*j*−1_ inherit the same symmetry with respect to ***δ***_*i*_ as in the two-step case, and the pairwise epistasis results provide a qualitative framework for understanding path-dependent epistasis along longer trajectories.

We simulated adaptive walks at two levels of phenotypic complexity (*n* = 10 and *n* = 100) and three distances to the optimum (*d* = 2.5, 0.25, and 0.07), with 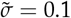 fixed throughout, corresponding to a mean fitness effect of beneficial mutations of approximately 1%. For each condition, we computed the generalized epistasis coefficient ℰ_*i,j*_ for a focal substitution *i* occurring at distance *d*, with *j* ranging from *i* − 16 to *i* + 16 (Figure 3). This parameter space spans Regime I below and above the critical threshold *x*\*, as well as Regime II. To facilitate biological interpretation, epistasis was normalized by the mean log-fitness gain of the focal substitution at distance *d*.

**Figure 3.**
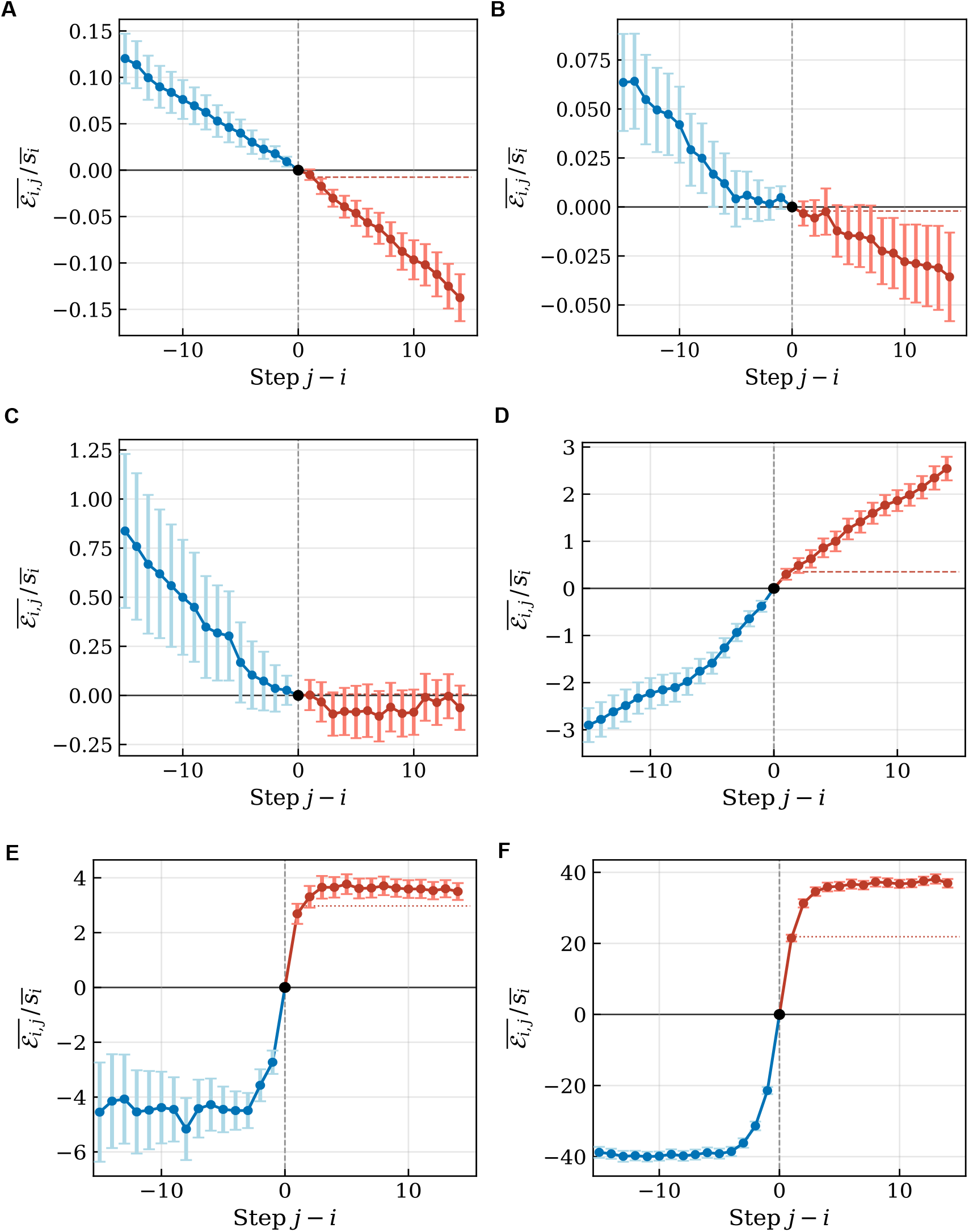
Contingency and entrenchment of a focal substitution along adaptive walks in Fisher’s geometric model. Columns correspond to phenotypic complexity (*n* = 10, panels **a**–**c**–**e**; *n* = 100, panels **b**–**d**–**f**); Rows correspond to decreasing distance to the optimum (*d* = 2.5, panels **a**,**b**; *d* = 0.25, panels **c**,**d**; *d* = 0.07, panels **e**,**f**), with 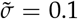 fixed throughout. The focal substitution is the 16th step of each walk, standardized to occur at distance *d* from the optimum (relative tolerance 10%). For each panel, 200 independent adaptive walks were simulated under the SSWM regime (beneficial mutations only, without weighting by fixation probability 2*s*). Blue circles show the mean generalized epistasis coefficient 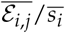 for *j < i* (contingency) and red circles for *j > i* (entrenchment), normalized by the mean log-fitness gain 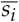 of the focal substitution. Error bars show ±2 standard errors across replicates. The black dot at *j* − *i* = 0 marks the focal substitution (ℰ _*i,i*_ = 0 by definition). The dashed line shows the Regime I asymptotic prediction for 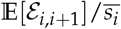 (equation 22); the dotted line shows the Regime II prediction (equation 23, displayed only when *y* ≥ 0.3). Dimensionless parameters for each panel: **(a)** *y* = 0.02, *x* = 0.063; **(b)** *y* = 0.02, *x* = 0.2; **(c)** *y* = 0.20, *x* = 0.63; **(d)** *y* = 0.20, *x* = 2; **(e)** *y* = 0.71, *x* = 2.26; **(f)** *y* = 0.71, *x* = 7.1. The critical threshold *x*\*≈ 0.612 separates diminishing returns (*x < x*\*) from entrenchment (*x > x*\*).

When the focal substitution arises far from the optimum and at low phenotypic complexity (*x < x*\*, Figure 3a), the predominant regime is one of diminishing returns. Successive substitutions are broadly aligned with a selection axis that varies little from step to step, so that the focal mutation would have been more beneficial had it arisen earlier, and becomes progressively easier to revert as later substitutions accumulate. Negative epistasis reflects the progressive flattening of the fitness landscape as the population approaches the optimum. Note that simulations incorporating fixation probability show qualitatively similar results, with the sign change occurring at a slightly larger *x* than *x*\* as weighting by 2*s* favors mutations more aligned with the optimum (Figure S1 and S4).

At higher complexity or closer to the optimum (*x > x*\*, Figure 3d), mutations carry a larger orthogonal component due to antagonistic pleiotropy: they improve fitness along the selection axis but simultaneously displace the phenotype along several other traits. Subsequent substitutions compensate these deleterious phenotypic components, making reversion of ***δ***_*i*_ increasingly costly: ***δ***_*i*_ is entrenched. Symmetrically, ***δ***_*i*_ is contingent on preceding mutations: it can fix only because it compensates the antagonistic pleiotropy accumulated by earlier substitutions. Without those preceding substitutions, ***δ***_*i*_ would have been deleterious or insufficiently beneficial to fix in the background **z**_*i*−*k*_.

A focal substitution arising near the transition *x* ≈ *x*\* (Fig-ure 3c) accumulates little epistasis with future substitutions: its reversion cost remains approximately equal to its original fitness benefit. Nevertheless, simulations show that it would have been significantly more beneficial in earlier backgrounds. This asymmetry reflects the directional nature of the adaptive walk: preceding substitutions occurred at larger distances from the optimum, where *x < x*\* and mutations pointed more directly toward the optimum, generating systematic alignment with ***δ***_*i*_and hence positive ℰ_*i,i*−*k*_even when local entrenchment is negligible.

Close to the optimum (Regime II, Figure 3e-f), entrenchment and contingency are both strong and systematic, and diminishing returns epistasis disappears entirely. The absolute magnitude of normalized epistasis is substantially larger in this regime, consistent with the analytical maximum at 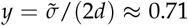 predicted by equation (23). For the experimentalist, this result carries a direct implication: substitutions observed in proteins that have evolved for a long time under purifying selection — and are therefore close to their phenotypic optimum — should systematically display strong signatures of both entrenchment and contingency.

### Contingency and entrenchment at mutation-selection-drift equilibrium

The results above assume exclusive fixation of beneficial mutations. Two features of the mutation-selection-drift equilibrium (MSDE) regime make it qualitatively distinct. First, weakly deleterious mutations can fix by genetic drift, so that the population is not confined to beneficial substitutions. Second, the stationary phenotypic distribution is much closer to the optimum than the regimes considered above. Under Gaussian fitness and the SSWM approximation, the mean log-fitness at stationarity is (Sella 2009; Tenaillon et al. 2007):

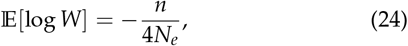

corresponding to a mean distance to the optimum of approximately

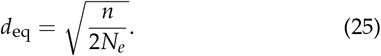

For 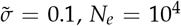 and *n* ranging from 5 to 100, this gives *d*_eq_ from 0.016 to 0.071, and 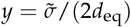 from 0.7 to 3.5 — values squarely in Regime II or at its boundary. Compared with the near-optimum Regime II considered above, the MSDE additionally involves frequent overshooting, with the selection axis shifting abruptly at each substitution as deleterious mutations push the phenotype away from the optimum in varying directions.

Despite this increased stochasticity, simulations confirm that both contingency and entrenchment are systematic and strong at MSDE (Figure 4). Entrenchment is large relative to drift, with |*N*_*e*_*ℰ*_*i,j*_| ≫ 1 for *n* ≥ 10. It establishes rapidly — within fewer than five substitutions — and then plateaus, indicating that the mutations immediately following the focal substitution are the ones that compensate its phenotypic effect, while later substitutions are orthogonal to it and contribute little additional entrenchment.

**Figure 4.**
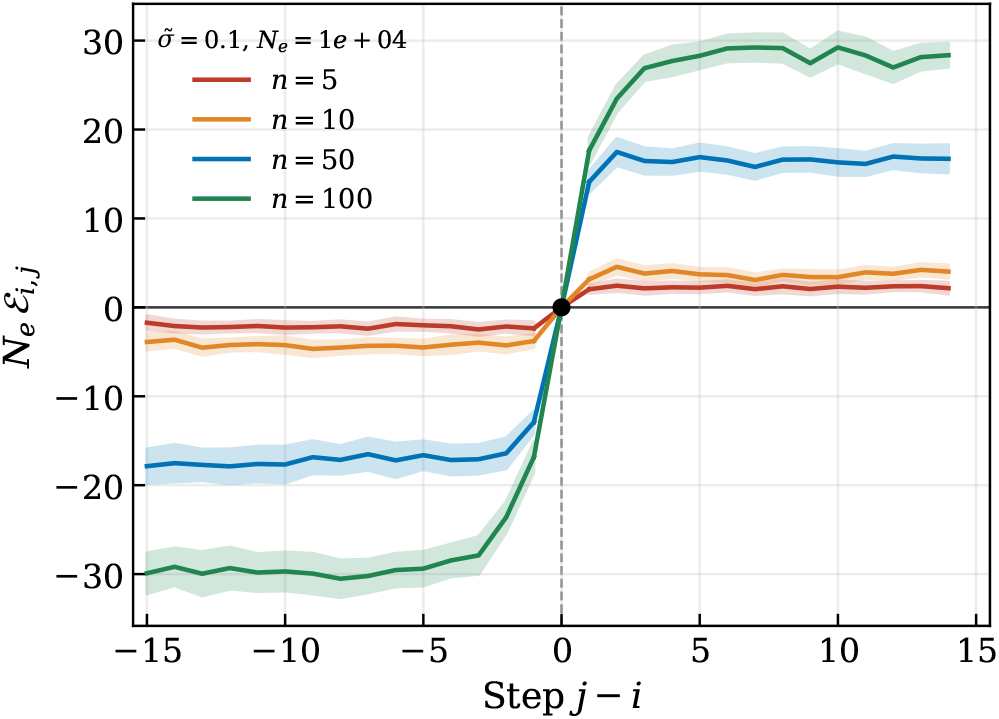
Path-dependent epistasis at mutation-selection-drift equilibrium. Contingency (*j < i*, negative values) and entrenchment (*j > i*, positive values) of a focal substitution along MSDE walks in Fisher’s geometric model, for four levels of phenotypic complexity (*n* = 5, 10, 50, 100), with 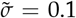 and *N*_*e*_ = 10^4^. Walks were initialized at the stationary distance 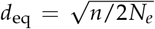 and simulated using Kimura fixation probabilities allowing both beneficial and deleterious mutations to fix (MCMC sampling). Curves show the mean *N*_*e*_ *ℰ*_*i,j*_ averaged over 50 replicates; shaded bands show ±2 standard errors.Both contingency and entrenchment increase strongly with *n*, consistent with the Regime II prediction *N*_*e*_ ℰ_*i,i*±1_ ~ *n* · *g*(*y*)^2^, where 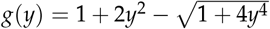 and 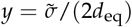.

Strikingly, both contingency and entrenchment increase markedly with phenotypic complexity *n*. This dependence can be understood quantitatively from the Regime II prediction. Since 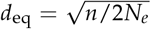 increases with *n*, larger *n* corresponds to smaller 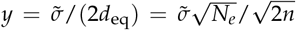. The expected pairwise epistasis in Regime II then scales as

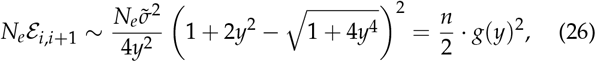

where 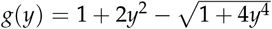. Two effects combine to amplify epistasis as *n* increases: the prefactor *n* grows directly, and *g*(*y*)^2^ also increases as *y* decreases toward the maximum of *g* at 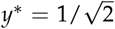. These two effects reinforce each other, producing the strong dependence on complexity observed in Figure 4.

From a biological perspective, these results demonstrate that contingency and entrenchment are not restricted to adaptive walks — they are a generic feature of protein-coding sequence evolution under purifying selection, arising whenever phenotypic complexity is non-trivial. Both phenomena are strong relative to genetic drift (|*N*_*e*_ *ℰ*_*i,j*_ | ≫ 1) and increase with phenotypic complexity. This implies that substitutions observed in proteins evolving under long-term purifying selection should systematically display signatures of historical contingency and entrenchment, with magnitude increasing with the number of traits under stabilizing selection.

## Discussion

Fisher’s geometric model (FGM) describes a smooth phenotypic landscape with stabilizing selection toward a single optimum. It was originally introduced to formalize the verbal argument that complex adaptations must proceed in small steps, because pleiotropy makes large mutations overwhelmingly likely to be deleterious (Fisher 1930). Despite its simplicity, nontrivial properties emerge from this abstract framework, and FGM has become a central *proof-of-concept* model in adaptation theory over the past three decades (Tenaillon 2014). Rather than tracing mutational effects to their underlying molecular basis, it identifies robust features of the adaptive process expected to hold across large classes of organisms. In this capacity, it has yielded analytical predictions for the distribution of fitness effects of new mutations, the size distribution of fixed substitutions, the rate of adaptation, and the cost of phenotypic complexity(Orr 2000, 2006; Martin and Lenormand 2006; Tenaillon *et al*. 2007). As emphasized by Servedio *et al*. (2014), the value of such models lies not in their quantitative realism but in their capacity to test the logical consistency of verbal hypotheses and to reveal predictions that would not follow from verbal reasoning alone.

FGM provides a generic scenario for the emergence of epistatic interactions from a nonlinear mapping of an additive, multidimensional phenotype onto fitness. Three partially overlapping mechanisms operate, with contrasting effects on the sign of epistasis. First, the nonlinear relationship between distance to the optimum and fitness produces diminishing returns: a given phenotypic displacement toward the optimum yields a larger fitness gain far from the optimum than close to it, a form of curvature ubiquitous in biology, from the sigmoidal relationship between protein stability and fraction folded to the saturating dependence of pathway flux on enzyme concentration (Kacser and Burns 1981). This mechanism generates negative epistasis. Second, when the population is close to the optimum, two individually beneficial mutations directed toward the optimum may jointly overshoot it generating negative epistasis. Third, when mutations affect more than one trait, a beneficial mutation generically displaces the phenotype in two distinct components: one toward the optimum, which is positively selected, and one perpendicular to it, which is penalized by the curvature of fitness isoclines. These two effects within a single mutation are antagonistic to one another: the mutation improves fitness along the main axis of selection at the cost of degrading the orthogonal traits, a mechanism Blanquart *et al*. (2014) connected to antagonistic pleiotropy. For a single beneficial mutation, the orthogonal component is by definition small enough relative to the component toward the optimum that the mutation remains beneficial. For a pair of successive mutations, however, the situation is more complex: the orthogonal displacement introduced by the first substitution generates a new selection pressure to compensate its collateral effects, biasing the distribution of the second substitution toward correcting this phenotypic damage. More formally, the first mutation reorients the selection axis experienced by the second: the direction of steepest fitness ascent is no longer aligned with the original selection axis, but shifted toward compensating the collateral effects of the first substitution. Because the second mutation is beneficial precisely because it compensates the side-effects of the first, its fitness gain is expected to be smaller, or even negative, when evaluated in a background lacking the first mutation — the signature of positive epistasis between successive substitutions. This work shows that this mechanism constitutes the geometric basis of contingency and entrenchment in FGM. Which of the three mechanisms predominates determines the sign of net epistasis along the walk. Beyond these pairwise considerations, epistasis among successive substitutions along an adaptive walk raises two additional difficulties: beneficial mutations are a strongly nonrandom subset of all mutations, and successive substitutions are not independent, so that natural selection is expected to bias both the prevalence and the type of epistatic interactions among the mutations that fix.

This work derives predictions for path-dependent epistasis among mutations that fix along adaptive trajectories in Fisher’s geometric model. First, we generalize the entrenchment and contingency coefficients introduced by Shah *et al*. (2015) to the FGM framework. These coefficients take a simple form and have a transparent geometric interpretation: entrenchment of a mutation is determined by its alignment with the subsequent adaptive path, and contingency by its alignment with the prior path.

Second, for adaptive trajectories that reach the optimum, entrenchment admits a geometric interpretation in terms of nested spheres, from which the probability that a beneficial mutation is entrenched can be derived analytically. This probability depends critically on the Fisher parameter 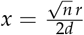, which jointly captures the effects of phenotypic complexity *n*, distance to the optimum *d*, and mutation size *r*, combining the curvature of fitness isoclines with the tendency of phenotypic complexity to misalign mutational vectors from the axis of selection.

Third, we derive analytical predictions for the sign and magnitude of the expected pairwise epistasis between two consecutive beneficial mutations, extending the results of Blanquart *et al*. (2014), which were restricted to the regime far from the optimum. The sign of mean epistasis depends on the same Fisher parameter *x*.

Fourth, simulations of longer adaptive walks confirm that the transition between diminishing returns epistasis and entrenchment is governed by the distance to the optimum and phenotypic complexity. Far from the optimum, the nonlinear fitness-distance relationship dominates and produces diminishing returns. As distance decreases or phenotypic complexity increases, antagonistic pleiotropy becomes the dominant mechanism: orthogonal components accumulate along the trajectory and are compensated by subsequent substitutions, generating entrenchment and contingency. Near the optimum, at mutation-selection-drift equilibrium, overshooting represents an extreme form of axis reorientation that further amplifies entrenchment and contingency.

Our results demonstrate that contingency, entrenchment, and diminishing returns epistasis all emerge as generic properties of adaptive trajectories in FGM under minimal assumptions: stabilizing selection toward a single optimum and pleiotropic mutations. A key conceptual contribution concerns the mechanistic basis of contingency and entrenchment, which both reflect positive epistasis between successive substitutions. It has been previously suggested that such effects primarily reflect specific epistasis, that is, in the context of protein evolution, direct physical interactions between particular residues (Starr and Thornton 2016). Our results show that this is not necessarily the case. Pervasive contingency and entrenchment arise in FGM entirely from nonspecific epistasis, through antagonistic pleiotropy: orthogonal phenotypic displacements accumulate across successive substitutions, each substitution progressively reorients the axis of selection, and subsequent mutations compensate for the orthogonal costs of earlier ones, making their reversion increasingly costly. Nonspecific epistasis is a generic consequence of any nonlinear mapping from a multidimensional phenotype to fitness and does not require any system-specific structural interaction. Our analytical framework further identifies the conditions under which each regime predominates and yields two quantitative and testable predictions. First, entrenchment increases with phenotypic complexity *n* at fixed mutation size, because more dimensions are available for orthogonal displacements to accumulate. Second, entrenchment increases as the population approaches the optimum: as the relative mutation size *r*/*d* grows, each substitution reorients the selection axis more substantially, amplifying the antagonistic pleiotropic costs of prior substitutions and the selective pressure to compensate for them.

Several limitations of the present analysis deserve mention. First, the analytical derivations weight all beneficial mutations equally, without accounting for their probability of fixation, which increases with selective coefficient. This bias may quantitatively shift predicted distributions of entrenchment and contingency, particularly near the optimum, although simulations suggest that the qualitative predictions are robust. Second, the analytical results are restricted to beneficial mutations; weakly deleterious mutations that fix by drift are excluded from the formal derivations, though their contribution is expected to be negligible far from the optimum where beneficial mutations dominate. Third, the model assumes isotropic mutational effects, a quadratic fitness function, and asexual reproduction. Extensions relaxing each of these assumptions exist (Waxman 2006; Martin and Lenormand 2006; Martin et al. 2007; Gros *et al*. 2009), but their quantitative consequences for entrenchment and contingency remain to be characterized.These limitations are inherent to any proof-of-concept framework and do not affect the qualitative predictions, which depend only on the geometry of stabilizing selection and the pleiotropic structure of mutations.

A similar shift in the sign of epistasis along adaptive trajectories had been reported in the NK model by (Draghi and Plotkin 2013). They found, much like our work, a predominance of antagonistic epistasis in the early steps of adaptation and of synergistic or sign epistasis later on. The mechanisms invoked are however fundamentally different. In the NK model, early antagonism arises through a regression-to-the-mean effect: large-effect mutations that fix early were selectively favored precisely because they formed favorable interactions with the ancestral background. Their fixation therefore partly undermines the benefits of mutations at interacting sites, which see their selective coefficients suppressed. Late synergistic epistasis, in contrast, reflects the finiteness of genotypic space: near a local fitness peak, the only remaining beneficial mutations are those unlocked by the epistatic effects of the last substitution, so that interacting pairs of substitutions become the only avenue for further adaptation. In our model, the same qualitative shift emerges from entirely different mechanisms. Early diminishing returns epistasis reflects the nonlinear fitness-distance relationship, as successive mutations aligned with the direction of selection experience progressively smaller fitness gains. Late positive epistasis reflects the accumulation of antagonistic pleiotropic costs, compensated by subsequent substitutions. Two additional features distinguish our results from those of Draghi and Plotkin (2013). First, in the NK model the sign change occurs necessarily near the end of the walk, whereas in our model the transition is governed by phenotypic complexity *n* and relative distance to the optimum *r*/*d*, and can occur well before the optimum is reached for sufficiently large *n*. Second, the NK model requires that deleterious mutations cannot fix, whereas positive epistasis persists in our model near the optimum even when weakly deleterious mutations are allowed to fix by drift, as in the mutation-selection-drift equilibrium studied by Shah *et al*. (2015).

These two regimes find support in contrasting bodies of experimental evidence. Evidence for diminishing returns epistasis comes primarily from experimental evolution studies in which bacterial populations adapt to a novel selective pressure from an initially maladapted state. Khan *et al*. (2011) measured epistasis among the first five substitutions to fix in an *E. coli* population of the LTEE after 2,000 generations, collectively responsible for a fitness gain of nearly 30%. Chou *et al*. (2011) documented a similar pattern during the adaptation of an engineered *Methylobacterium extorquens* strain to growth on methanol, with an average fitness increase of 67% over 600 generations. In both cases, successive beneficial mutations interact antagonistically, consistent with the regime far from the optimum where diminishing returns dominate. Evidence for entrenchment and contingency comes from a different class of studies, focused on proteins evolving under long-term purifying selection near their functional optimum. Shah *et al*. (2015) documented pervasive entrenchment in model of proteins evolving under stabilizing selection over hundreds of millions of years. This regime likely characterizes the evolution of most genes underlying core cellular functions (Starr *et al*. 2018; Park *et al*. 2022), where populations remain close to a fitness optimum and weakly deleterious mutations fix by drift before being compensated.

Fisher’s geometric model unifies these two apparently distinct phenomena under a single mechanistic framework. Diminishing returns far from the optimum and entrenchment near it are not the products of different evolutionary processes but emerge from the same underlying geometry. The transition between these two regimes is controlled by a single composite parameter combining phenotypic complexity *n* and relative distance to the optimum *r*/*d*, which determines whether alignment with selection or orthogonal pleiotropy dominates the epistatic structure of adaptive trajectories. More broadly, the Fisher parameter *x* provides a single axis along which disparate empirical observations of diminishing returns and entrenchment can be ordered, suggesting that these phenomena reflect a common continuum whose position can in principle be inferred from estimates of phenotypic complexity, mutation size, and degree of maladaptation

## Data availability

This study is entirely theoretical. All analytical results are derived in the Appendices. Code used to generate all simulation results and figures is available at https://github.com/[repository].

