## Appendix for "From Diminishing Returns to Entrenchment: A Unifying Theory of Epistasis along Adaptive Walks Revealed by Fisher’s Geometric Model"

May 5, 2026

**5 Supplementary figures** **15**

---

### 1 Model and notation

#### 1.1 Fisher's geometric model

We work in $\mathbb{R}^n$, where $n$ denotes the number of phenotypic dimensions under stabilizing selection. The phenotypic optimum is placed at the origin. A phenotype is a vector $\mathbf{z} \in \mathbb{R}^n$; its fitness is

$$W(z) = e^{-\|\mathbf{z}\|^2/2}, \qquad \text{so that} \quad w(\mathbf{z}) := \log W(\mathbf{z}) = -\frac{\|\mathbf{z}\|^2}{2}.$$

A population is represented by its mean phenotype $\mathbf{z}_i$ at substitution step $i$, at distance $d_i = \|\mathbf{z}_i\|$ from the optimum. Each mutation is a random vector

$$\delta \sim \mathcal{N}(0, \sigma^2 I_n),$$

drawn independently at each step. Under the strong-selection weak-mutation (SSWM) regime, the population is monomorphic at all times and adaptation proceeds through the sequential fixation of new mutations. After fixation of $\delta_i$:

$$\mathbf{z}_i = \mathbf{z}_{i-1} + \delta_i.$$

The log-fitness change induced by mutation $\delta$ from position $z$ is

$$w(\mathbf{z} + \delta) - w(\mathbf{z}) = -\langle \mathbf{z}, \delta \rangle - \frac{\|\delta\|^2}{2}.$$

#### 1.2 Geometric parameterisation

For a mutation $\delta_i$ from position $\mathbf{z}_{i-1}$ with $d_{i-1} = \|\mathbf{z}_{i-1}\|$, we define:

- $r_i = \|\delta_i\|$, the Euclidean norm of the mutational vector;
- $\theta_i = \angle(\delta_i, \mathbf{z}_{i-1}) \in [0, \pi]$, the angle between $\delta_i$ and $\mathbf{z}_{i-1}$, defined so that $\langle \mathbf{z}_{i-1}, \delta_i \rangle = r_i d_{i-1} \cos\theta_i$;
- $\tilde{\sigma} = \sigma\sqrt{n}$, the typical mutational norm (since $\mathbb{E}[\|\delta\|] \approx \sigma\sqrt{n}$ for large $n$);
- $x = \dfrac{n\sigma}{2d_{i-1}}$, the dimensionless Fisher parameter.
- $y = \dfrac{\tilde{\sigma}}{2d_{i-1}}$, half the approximate relative distance to optimum.

With this parameterisation:

$$d_i^2 = d_{i-1}^2 + 2r_i d_{i-1}\cos\theta_i + r_i^2.$$

**Beneficial condition.** A mutation is beneficial ($d_i < d_{i-1}$) if and only if

$$\cos\theta_i < -\frac{r_i}{2d_{i-1}},$$

which requires $\theta_i \in (\pi/2, \pi]$ and $r_i < 2d_{i-1}$.

The following quantities appear repeatedly in the integration limits:

- $\gamma(r_i) = \arccos\left(-\dfrac{r_i}{2d_{i-1}}\right) \in (\pi/2, \pi)$, the minimum angle for beneficiality at norm $r_i$;
- $a(\theta_i) = -2d_{i-1}\cos\theta_i > 0$ for $\theta_i \in (\pi/2, \pi)$, the maximum norm for beneficiality at angle $\theta_i$.

These two quantities are related by $a(\gamma(r)) = r$ and $\gamma(a(\theta)) = \theta$.

#### 1.3 Concentration of the mutation angle in high dimension

We justify the concentration of the angle $\theta$ between a mutation and the current phenotype around $\pi/2$ as $n \to \infty$, for isotropic mutations. This result applies in particular to $\delta \sim \mathcal{N}(0, \sigma^2 I_n)$.

In hyperspherical coordinates, the marginal density of $\theta \in [0, \pi]$ for an isotropic distribution on $\mathbb{S}^{n-1}$ is:

$$f(\theta) = \frac{\sin^{n-2}\theta}{\displaystyle\int_0^\pi \sin^{n-2}\theta\, d\theta}.$$

The denominator is a Wallis integral; a classical result gives:

$$\int_0^\pi \sin^{n-2}\theta\, d\theta \underset{n\to\infty}{\sim} \sqrt{\frac{2\pi}{n}}.$$

Hence $f(\theta) \sim \sqrt{n/(2\pi)}\, \sin^{n-2}\theta$, which is sharply concentrated around $\theta = \pi/2$ where $\sin\theta = 1$.

We introduce the rescaled variables:

$$X_n = \sqrt{n}\left(\theta - \frac{\pi}{2}\right), \qquad Y_n = \sqrt{n}\cos\theta.$$

The substitution $u = \sqrt{n}(\theta - \pi/2)$ for $X_n$ and $u = \sqrt{n}\cos\theta$ for $Y_n$ yields the densities:

$$f_{X_n}(u) = \frac{1}{\sqrt{2\pi}}\cos^{n-2}\left(\frac{u}{\sqrt{n}}\right) \cdot \mathbf{1}_{\left[-\frac{\sqrt{n}\pi}{2}, \frac{\sqrt{n}\pi}{2}\right]}(u),$$

$$f_{Y_n}(u) = \frac{1}{\sqrt{2\pi}}\left(1 - \frac{u^2}{n}\right)^{(n-3)/2} \cdot \mathbf{1}_{[-\sqrt{n}, \sqrt{n}]}(u).$$

Both densities converge pointwise to $e^{-u^2/2}/\sqrt{2\pi}$ as $n \to \infty$. By Scheffé's theorem, this pointwise convergence of densities implies convergence in distribution:

$$\sqrt{n}\cos\theta \xrightarrow[n\to\infty]{\mathcal{L}} \mathcal{N}(0,1), \qquad \sqrt{n}\left(\theta - \frac{\pi}{2}\right) \xrightarrow[n\to\infty]{\mathcal{L}} \mathcal{N}(0,1).$$

Equivalently, for large $n$:

$$\cos\theta \approx \mathcal{N}\left(0, \frac{1}{n}\right), \qquad \theta \approx \mathcal{N}\left(\frac{\pi}{2}, \frac{1}{n}\right).$$

This equatorial concentration is a classical manifestation of the curse of dimensionality: in high dimension, any fixed direction becomes nearly orthogonal to a uniformly random vector on $\mathbb{S}^{n-1}$.

#### 1.4 Density of beneficial mutations

Consider now a mutation $\delta_i$ arising from background $z_{i-1}$, with $d = d_{i-1} = \|z_{i-1}\|$.Since $\delta_i \sim \mathcal{N}(0, \sigma^2 I_n)$ is isotropic, its density depends only on $r_i = \|\delta_i\|$ and $\theta_i = \angle(\delta_i, z_{i-1})$. The beneficial condition $\cos\theta_i < -r_i/(2d)$ restricts the support to the region $\theta_i \in (\pi/2, \pi]$, $r_i \in (0, 2d)$.

In hyperspherical coordinates in $\mathbb{R}^n$, the density of the mutational vector conditional on being beneficial is:

$$f_B(r,\theta \mid d) = \frac{1}{K_B}\, r^{n-1} e^{-r^2/(2\sigma^2)} \sin^{n-2}\theta \; \mathbf{1}\Big\{\cos\theta \le -\frac{r}{2d},\ r < 2d\Big\}\,, \tag{1}$$

where the normalisation constant is obtained by integrating over the ball of radius $d$ centered at the optimum:

$$K_B = \int_0^{2d} \int_{\gamma(r)}^{\pi} r^{n-1}\, e^{-r^2/(2\sigma^2)}\, \sin^{n-2}\theta\; d\theta, dr \tag{2}$$

with $\gamma(r) = \arccos(-r/2d)$ the maximum beneficial angle at norm r.

##### 1.5 Asymptotic regimes

We consider two asymptotic regimes, characterised respectively by $x = O(1)$ and $r/d = O(1)$, as $n \to \infty$.

###### 1.5.1 Regime I: $x = O(1)$

This regime corresponds to a population whose distance to the optimum grows as $\sqrt{n}$ with dimensionality, while the typical mutational norm $\tilde{\sigma}$ remains fixed.The Fisher parameter $x = n\sigma/(2d)$ is then held at a fixed finite value.

$$d = D\sqrt{n}, \quad D > 0 \text{ fixed}; \qquad \sigma = \frac{\tilde{\sigma}}{\sqrt{n}}, \quad \tilde{\sigma} > 0 \text{ fixed}; \qquad x = \frac{\tilde{\sigma}}{2D} = O(1), \quad n \to \infty. \tag{3}$$

Equivalently, at fixed distance $d$, this regime arises when $\tilde{\sigma}$ is sufficiently small that $x = \tilde{\sigma}\sqrt{n}/(2d) = O(1)$.

A classical result of FGM shows that in this regime, the probability for a mutation to be beneficial satisfies

$$P_{\text{ben}} \underset{n\to\infty}{\sim} 1 - \Phi(x), \tag{4}$$

where $\Phi$ is the standard normal CDF. The parameter $x$ thus controls the fraction of beneficial mutations: large $x$ corresponds to a population close to the optimum (or high complexity), where beneficial mutations are rare.

Three angular integrals arise repeatedly. For $r = \tilde{\sigma}\, t$, the substitution $u = -\cos\theta$ then $v = \sqrt{n}\, u$ ($G_0$ and $G_2$), together with dominated convergence ($(1 - v^2/n)^{(n-3)/2} \to e^{-v^2/2}$), gives:

$$G_0(\tilde{\sigma}t) := \int_{\gamma(\tilde{\sigma}t)}^{\pi} \sin^{n-2}\theta\, d\theta \underset{n\to\infty}{\sim} \frac{1}{\sqrt{n}} \int_{xt}^{\infty} e^{-v^2/2}\, dv, \tag{5}$$

$$G_1(\tilde{\sigma}t) := \int_{\gamma(\tilde{\sigma}t)}^{\pi} \sin^{n-2}\theta \cos\theta\, d\theta = -\frac{1}{n-1}\left(1 - \frac{x^2t^2}{n}\right)^{(n-1)/2} \underset{n\to\infty}{\sim} -\frac{1}{n}\, e^{-x^2t^2/2}, \tag{6}$$

$$G_2(\tilde{\sigma}t) := \int_{\gamma(\tilde{\sigma}t)}^{\pi} \sin^{n-2}\theta \cos^2\theta\, d\theta \underset{n\to\infty}{\sim} \frac{1}{n^{3/2}} \int_{xt}^{\infty} v^2\, e^{-v^2/2}\, dv. \tag{7}$$

Evaluating at $t = 1$:

$$G_0(\tilde{\sigma}) \sim \frac{\sqrt{2\pi}}{\sqrt{n}}\,(1 - \Phi(x)), \qquad G_1(\tilde{\sigma}) \sim -\frac{1}{n}\, e^{-x^2/2}, \qquad G_2(\tilde{\sigma}) \sim \frac{1}{n^{3/2}} \int_x^{\infty} v^2\, e^{-v^2/2}\, dv.$$

##### 1.5.2 Regime II: $y = \tilde{\sigma}/2d = O(1)$

$$d = O(1), \quad \sigma = \frac{\tilde{\sigma}}{\sqrt{n}}, \quad \tilde{\sigma} > 0 \text{ fixed}, \qquad n \to \infty, \tag{8}$$

so that $x = n\sigma/(2d) = \tilde{\sigma}\sqrt{n}/(2d) \to \infty$. This regime corresponds to a population close to the optimum relative to the typical mutational norm: beneficial mutations become rare and their distribution concentrates near the boundary of the beneficial region.

#### 1.6 Special functions

We denote by $\varphi$ and $\Phi$ the standard normal density and distribution function:

$$\varphi(x) = \frac{e^{-x^2/2}}{\sqrt{2\pi}}, \qquad \Phi(x) = \int_{-\infty}^{x} \varphi(t)\, dt.$$

**Inverse Mills ratio (hazard rate).**

$$\Lambda(x) := \frac{\varphi(x)}{1 - \Phi(x)}.$$

**Key properties**: $\Lambda(0) = \sqrt{2/\pi}$; $\Lambda'(x) = \Lambda(x)(\Lambda(x) - x) > 0$ (strictly increasing). The strict positivity of $\Lambda'$ follows from inequality $1 - \Phi(x) < \varphi(x)/x$ for all $x > 0$,

Moreover $\Lambda'(x) \leq 1$ for all $x > 0$, which is a consequence of the Cauchy–Schwarz inequality applied to functions 1 and $t$ under the measure $\varphi(t)\, dt$ on $(x, +\infty)$:

$$\varphi(x)^2 = \left(\int_x^\infty t\, \varphi(t)\, dt\right)^2 \leq \underbrace{\int_x^\infty \varphi(t)\, dt}_{F(x)} \cdot \underbrace{\int_x^\infty t^2\, \varphi(t)\, dt}_{x\varphi(x) + F(x)},$$

The complementary CDF of the standard normal admits the classical asymptotic expansion

$$1 - \Phi(x) = \frac{\phi(x)}{x}\left(1 - \frac{1}{x^2} + \frac{3}{x^4} - \frac{15}{x^6} + \cdots\right) = \frac{\phi(x)}{x} \sum_{k=0}^{\infty} \frac{(-1)^k (2k-1)!!}{x^{2k}}, \tag{9}$$

as $x \to +\infty$, where $(2k-1)!! = 1 \cdot 3 \cdots (2k-1)$ denotes the double factorial. Setting $u = 1/x^2 - 3/x^4 + \cdots$, the inverse Mills ratio $\Lambda(x) = \phi(x)/(1 - \Phi(x))$ satisfies

$$\Lambda(x) = \frac{x}{1-u} = x(1 + u + u^2 + \cdots) = x\left(1 + \frac{1}{x^2} - \frac{2}{x^4} + \cdots\right), \tag{10}$$

hence

$$\Lambda(x) \sim x + \frac{1}{x} - \frac{2}{x^3} + O(x^{-5}), \qquad x \to +\infty. \tag{11}$$

### 2 Generalized epistasis in Fisher's geometric model

#### 2.1 Pairwise epistasis

Standard pairwise epistasis between mutations $\delta_1$ and $\delta_2$ is defined as the deviation of the log-fitness effect of the double mutant from the sum of the single-mutant effects:

$$\varepsilon_{1,2} := \big[w(z_0 + \delta_1 + \delta_2) - w(z_0)\big] - \big[w(z_0 + \delta_1) - w(z_0)\big] - \big[w(z_0 + \delta_2) - w(z_0)\big]. \tag{12}$$

As emphasized by Shah et al. (2015), this is equivalently written as the difference between the fitness effect of $\delta_2$ in the background carrying $\delta_1$ and its effect in the naive background:

$$\varepsilon_{1,2} = \big[w(z_0 + \delta_1 + \delta_2) - w(z_0 + \delta_1)\big] - \big[w(z_0 + \delta_2) - w(z_0)\big]. \tag{13}$$

This formulation makes the interpretation in terms of substitution order explicit: $\varepsilon_{1,2}$ measures how much the fitness effect of $\delta_2$ changes when $\delta_1$ is already present in the background, or equivalently, what would have been the fitness effect of $\delta_2$ if the order of the two substitutions were reversed. Expanding using $w(z) = -\|z\|^2/2$ we immediatly get:

$$\varepsilon_{1,2} = -\langle \delta_1, \delta_2 \rangle.$$

This interpretation suggests a natural generalization to longer evolutionary trajectories, originally proposed by Shah et al. (2015).

##### 2.2 Definition of generalized epistasis following Shah et al. (2015)

Consider a trajectory fixing substitutions $\delta_1, \delta_2, \ldots$ sequentially, with $z_i = z_{i-1} + \delta_i$. For any pair of substitutions $i$ and $j$, one can ask how much larger the fitness effect of substitution $\delta_i$ would have been under the alternative trajectory in which $\delta_i$ is removed from its actual position $i$ and instead inserted at position $j$. The *generalized epistasis coefficient* between substitution $i$ and position $j$ is defined as:

$$\mathcal{E}_{i,j} := \begin{cases} \big[w(z_{j-1} + \delta_i) - w(z_{j-1})\big] - \big[w(z_i) - w(z_{i-1})\big] & \text{if } i > j, \\ \big[w(z_j) - w(z_j - \delta_i)\big] - \big[w(z_i) - w(z_{i-1})\big] & \text{if } i < j. \end{cases} \tag{14}$$

It is straightforward to verify that $\mathcal{E}_{i,i+1}$ reduces to the standard pairwise epistasis $\varepsilon_{i,i+1}$ between two consecutive substitutions.

**Biological interpretation.** The two cases in (14) capture distinct biological phenomena.

**Case $i > j$ — contingency.** $\mathcal{E}_{i,j}$ compares the fitness effect of $\delta_i$ in background $z_{j-1}$ (had it occurred earlier, at step $j$) to its actual effect in $z_{i-1}$ (where it really occurred). A positive value, $\mathcal{E}_{i,j} > 0$, means that $\delta_i$ would have been more beneficial at step $j$ than it actually was at step $i$: the intervening substitutions $\delta_j, \ldots, \delta_{i-1}$ have made $\delta_i$ *less* beneficial, a form of diminishing returns. Conversely, $\mathcal{E}_{i,j} < 0$ means that $\delta_i$ would have been less beneficial — or even deleterious — in the earlier background $z_{j-1}$, but became beneficial only thanks to the preceding substitutions $\delta_j, \ldots, \delta_{i-1}$: substitution $\delta_i$ is then said to be *contingent* on those preceding substitutions.

**Case $i < j$ — entrenchment.** $\mathcal{E}_{i,j}$ compares the fitness cost of reverting $\delta_i$ in the later background $z_j$ (i.e. the fitness gain of going from $z_j - \delta_i$ to $z_j$) to the original fitness benefit of $\delta_i$ at fixation. A positive value, $\mathcal{E}_{i,j} > 0$, means that reverting $\delta_i$ in context $z_j$ is more deleterious than it would have been immediately after fixation: the subsequent substitutions $\delta_{i+1}, \ldots, \delta_j$ have made $\delta_i$ harder to remove, and substitution $\delta_i$ is said to be *entrenched* by those later substitutions. Conversely, $\mathcal{E}_{i,j} < 0$ means that $\delta_i$ is easier to revert after accumulation of $\delta_{i+1}, \ldots, \delta_j$, indicating that later substitutions have compensated for or reduced the fitness contribution of $\delta_i$.

##### 2.3 Closed form in Fisher's geometric model

Under the log-fitness $w(z) = -\|z\|^2/2$, a direct expansion shows that the $\|\delta_i\|^2$ terms cancel in both cases of (14), yielding:

$$\mathcal{E}_{i,j} = \begin{cases} \langle \delta_i,\ z_{i-1} - z_{j-1} \rangle = \sum_{k=j}^{i-1} \langle \delta_i,\, \delta_k \rangle & \text{if } i \geq j, \\ -\langle \delta_i,\ z_j - z_i \rangle = -\sum_{k=i+1}^{j} \langle \delta_i,\, \delta_k \rangle & \text{if } i < j. \end{cases} \tag{15}$$

In both cases $\mathcal{E}_{i,i} = 0$, and epistasis decomposes as a sum of inner products between $\delta_i$ and the intervening mutational vectors, independently of the absolute phenotypic position.Moreover as expected $\mathcal{E}_{i,i+1}$ is equivalent to paiwise epistasis

For consecutive substitutions, equation (15) gives:

$$\mathcal{E}_{i,i-1} = \langle \delta_i,\, \delta_{i-1} \rangle, \qquad \mathcal{E}_{i,i+1} = -\langle \delta_i,\, \delta_{i+1} \rangle. \tag{16}$$

These two quantities satisfy $\mathcal{E}_{i+1,i} = -\mathcal{E}_{i,i+1}$, which has a transparent interpretation: $\mathcal{E}_{i+1,i}$ measures the fitness gain of $\delta_{i+1}$ in background $z_{i-1}$ relative to its gain in $z_i$, which is precisely the negative of the standard pairwise epistasis $-\langle \delta_i,\, \delta_{i+1} \rangle$, the latter comparing the fitness effect of $\delta_{i+1}$ in background $z_i$ to its effect in $z_{i-1}$.

The sign of $\mathcal{E}_{i,j}$ determines the nature of the epistatic interaction. For $i > j$: $\mathcal{E}_{i,j} > 0$ means $\delta_i$ would have been more beneficial at the earlier position $j$, while $\mathcal{E}_{i,j} < 0$ indicates contingency - $\delta_i$ benefited from the intervening substitutions to become beneficial.

For $i < j$: $\mathcal{E}_{i,j} > 0$ means the reversion of $\delta_i$ becomes increasingly costly as substitutions $\delta_{i+1}, \ldots, \delta_j$ accumulate (entrenchment), while $\mathcal{E}_{i,j} < 0$ indicates that later substitutions have reduced the fitness contribution of $\delta_i$.

#### 3 Generalized epistasis at the optimum: $\mathcal{E}_{1,\infty}$

We consider the limiting case in which, starting from $z_0$, the adaptive walk eventually reaches the phenotypic optimum, so that $z_\infty = 0$. The generalized epistasis coefficient between the first substitution $\delta$ and the optimum is then, by (15):

$$\mathcal{E}_{1,\infty} = \langle \delta,\ z_0 - z_\infty \rangle = \langle \delta,\, z_0 + \delta - z_\infty - \delta \rangle = \langle \delta,\, z_0 + \delta \rangle.$$

We therefore study

$$S := \langle \delta,\, \delta + z_0 \rangle = \underbrace{\|\delta\|^2}_{=:A} + \underbrace{\langle \delta,\, z_0 \rangle}_{=:B}, \tag{17}$$

where $\delta$ is the first beneficial mutation from $z_0$, and we write $d = \|z_0\|$ throughout this section.

##### 3.1 Geometric interpretation

We introduce the midpoint $\mathbf{c} := z_0/2$ and rewrite $\mathcal{E}_{1,\infty} = \|z_1\|^2 - \langle z_0,\, z_1 \rangle$ in terms of the distance from $z_1$ to $\mathbf{c}$:

$$\|z_1 - \mathbf{c}\|^2 = \|z_1\|^2 - \langle z_0,\, z_1 \rangle + \frac{d_0^2}{4} = \mathcal{E}_{1,\infty} + \frac{d_0^2}{4},$$

where $d_0 = \|z_0\|$. Therefore:

$$\mathcal{E}_{1,\infty} = \|z_1 - \mathbf{c}\|^2 - \frac{d_0^2}{4} \tag{18}$$

The level sets of $\mathcal{E}_{1,\infty}$ are spheres centered at $\mathbf{c} = z_0/2$. In particular, $\delta_1$ is entrenched ($\mathcal{E}_{1,\infty} > 0$) if and only if $z_1$ lies outside the sphere of center $\mathbf{c}$ and radius $d_0/2$ — the sphere passing through both the origin and $z_0$ — and contingent ($\mathcal{E}_{1,\infty} < 0$) if and only if $z_1$ lies inside.

#### 3.2 Expectation of $S$ in Regime I

We derive $\mathbb{E}[S] = \mathbb{E}_{f_B}[\mathcal{E}_{1,\infty}]$ for large $n$ in Regime I, where $x = n\sigma/(2d) = O(1)$. By linearity:

$$\mathbb{E}[S] = \mathbb{E}[A] + \mathbb{E}[B].$$

##### 3.2.1 Term $A$: $\mathbb{E}[\|\delta\|^2]$

With the substitution $r = \tilde{\sigma}\, t$, we immediately have:

$$\mathbb{E}[\|\delta\|^2] = \tilde{\sigma}^2 \frac{\displaystyle\int_0^{\sqrt{n}/x} t^{n+1}\, e^{-nt^2/2}\, G_0(\tilde{\sigma}t)\, dt}{\displaystyle\int_0^{\sqrt{n}/x} t^{n-1}\, e^{-nt^2/2}\, G_0(\tilde{\sigma}t)\, dt}.$$

The Laplace method gives concentration of both integrals around $t^* = \operatorname{argmax}\big(\log t - t^2/2\big) = 1$. At $t = 1$, the factor $G_0(\tilde{\sigma}t)$ and laplace leading termes cancels between numerator and denominator hence:

$$\mathbb{E}[\|\delta\|^2] \underset{n\to\infty}{\sim} \tilde{\sigma}^2 = n\sigma^2. \tag{19}$$

##### 3.2.2 Term $B$: $\mathbb{E}[\langle\delta,\, z_0\rangle]$

Since $\langle\delta,\, z_0\rangle = rd\cos\theta$, similar change of variable $r = \tilde{\sigma}t$ and $u = \sin\theta$ gives:

$$\mathbb{E}[\langle\delta,\, z_0\rangle] = \frac{-\tilde{\sigma}\, d}{n-1} \frac{\displaystyle\int_0^{\sqrt{n}/x} t^{n}\, e^{-nt^2/2} \left(1 - \frac{x^2t^2}{n}\right)^{(n-1)/2} dt}{\displaystyle\int_0^{\sqrt{n}/x} t^{n-1}\, e^{-nt^2/2}\, G_0(\tilde{\sigma}t)\, dt}.$$

The Laplace method gives concentration of both integrals around $t^* = \operatorname{argmax}(\log t - t^2/2) = 1$. At $t = 1$, using $\big(1 - x^2t^2/n\big)^{(n-1)/2} \sim e^{-x^2t^2/2}$ and the expression (5) for $G_0$:

$$\mathbb{E}[\langle\delta,\, z_0\rangle] \underset{n\to\infty}{\sim} \frac{-\tilde{\sigma}\, d}{\sqrt{n}} \frac{e^{-x^2/2}}{\displaystyle\int_x^{\infty} e^{-v^2/2}\, dv}.$$

Using definition of $\Lambda(x)$:

$$\mathbb{E}[\langle\delta,\, z_0\rangle] \underset{n\to\infty}{\sim} -\frac{\tilde{\sigma}\, d}{\sqrt{n}} \Lambda(x) = -d\sigma\, \Lambda(x). \tag{20}$$

##### 3.2.3 Assembly and sign change

Combining (19) and (20), and using $n\sigma^2 = 2xd\sigma$ from the definition $x = n\sigma/(2d)$:

$$\mathbb{E}[\mathcal{E}_{1,\infty}] \underset{n\to\infty}{\sim} d\sigma\big(2x - \Lambda(x)\big). \tag{21}$$

The sign of $\mathbb{E}[S]$ is determined by the competition between $2x$ and $\Lambda(x)$. Since $\Lambda(x) - 2x$ is strictly decreasing (section 1.6), with $\Lambda(0) = \sqrt{2/\pi}$ and $\Lambda(x) \sim x$ as $x \to \infty$, the function $x \mapsto \Lambda(x) - 2x$ is continuous and changes sign exactly once. The equation $\Lambda(x^*) = 2x^*$ has the unique solution

$$x^* \approx 0.612, \tag{22}$$

giving:

$$\mathbb{E}[S] < 0 \ \text{ for } \ x < x^*, \qquad \mathbb{E}[S] > 0 \ \text{ for } \ x > x^*.$$

##### 3.3 Variance of $S$ in Regime I

By the same substitution $r = \tilde{\sigma}\, t$ and Laplace concentration at $t^* = 1$, the three second-order moments are:

$$\mathbb{E}[\|\delta\|^4] \underset{n\to\infty}{\sim} \tilde{\sigma}^4 = 4x^2\, d^2\sigma^2, \tag{23}$$

$$\mathbb{E}\big[\langle\delta,\, z_0\rangle^2\big] = d^2\tilde{\sigma}^2\, \frac{\displaystyle\int_0^{\sqrt{n}/x} t^{n+1}\, e^{-nt^2/2}\, G_2(\tilde{\sigma}t)\, dt}{\displaystyle\int_0^{\sqrt{n}/x} t^{n-1}\, e^{-nt^2/2}\, G_0(\tilde{\sigma}t)\, dt} \underset{n\to\infty}{\sim} d^2\sigma^2\, (1 + x\Lambda(x)), \tag{24}$$

$$\mathbb{E}\big[\|\delta\|^2\, \langle\delta,\, z_0\rangle\big] = \frac{-\tilde{\sigma}^3 d}{n-1}\, \frac{\displaystyle\int_0^{\sqrt{n}/x} t^{n+2}\, e^{-nt^2/2} \left(1 - \frac{x^2t^2}{n}\right)^{(n-1)/2} dt}{\displaystyle\int_0^{\sqrt{n}/x} t^{n-1}\, e^{-nt^2/2}\, G_0(\tilde{\sigma}t)\, dt} \underset{n\to\infty}{\sim} -2x\, d^2\sigma^2\, \Lambda(x). \tag{25}$$

Expanding $\mathrm{Var}[S] = \mathbb{E}[(A+B)^2] - (\mathbb{E}[S])^2$ and substituting (23)–(25) with (21):

$$\mathrm{Var}[S] \underset{n\to\infty}{\sim} d^2\sigma^2\, \big(1 + x\Lambda(x) - \Lambda(x)^2\big) \tag{26}$$

**Large-$x$ asymptotics of $\mathrm{Var}[\mathcal{E}_{1,\infty}]$.** From equation (26), the variance is $\mathrm{Var}[\mathcal{E}_{1,\infty}] = d^2\sigma^2 f(x)$ with $f(x) = 1 + x\Lambda(x) - \Lambda(x)^2$. Writing $\epsilon(x) := \Lambda(x) - x = 1/x - 2/x^3 + O(x^{-5})$ for large $x$, a direct expansion gives

$$\begin{aligned} f(x) &= 1 - x\epsilon(x) - \epsilon(x)^2 \\ &= 1 - \left(1 - \frac{2}{x^2}\right) - \frac{1}{x^2} + O(x^{-4}) \\ &= \frac{1}{x^2} + O(x^{-4}). \end{aligned} \tag{27}$$

Therefore:

$$\mathrm{Var}[\mathcal{E}_{1,\infty}] \underset{x\to\infty}{\sim} \frac{d^2\sigma^2}{x^2}, \tag{28}$$

and the coefficient of variation satisfies $CV \sim 1/x^2 \to 0$: entrenchment becomes increasingly deterministic as $x$ increases.

#### 4 Epistasis between consecutive beneficial mutations

We now derive the expected inner product $\mathbb{E}[\langle \delta_i,\, \delta_{i+1}\rangle]$ between two consecutive beneficial mutations, which by (16) equals $-\mathcal{E}_{i,i+1}$.

##### 4.1 Local frame after fixation of $\delta_i$

After fixation of $\delta_i$ from $z_{i-1}$, the population sits at $z_i = z_{i-1} + \delta_i$, at distance

$$d_i = \|z_i\| = \sqrt{d_{i-1}^2 + 2r_i d_{i-1}\cos\theta_i + r_i^2}$$

from the optimum. Without loss of generality, we place $z_{i-1} = d_{i-1}\, e_1$ and $\delta_i = r_i(\cos\theta_i,\, \sin\theta_i,\, 0, \ldots, 0)$ in the plane spanned by $e_1$ and $e_2$. The new position is then

$$z_i = (d_{i-1} + r_i\cos\theta_i,\ r_i\sin\theta_i,\ 0, \ldots, 0).$$

We introduce a local orthonormal frame adapted to $z_i$. The radial direction from the optimum is $e_1' = z_i/d_i$. In the plane $(z_{i-1}, \delta_i)$, we define the unit vector orthogonal to $e_1'$:

$$u = \frac{1}{d_i}\big(-r_i\sin\theta_i,\ d_{i-1} + r_i\cos\theta_i,\ 0, \ldots, 0\big),$$

and let $\{v_3, \ldots, v_n\}$ be an arbitrary orthonormal basis of $(e_1', u)^\perp$. The next mutation $\delta_{i+1}$ is parameterised as

$$\delta_{i+1} = r_{i+1}\cos\varphi\ e_1' + r_{i+1}\sin\varphi\cos\psi\ u + r_{i+1}\sin\varphi\sin\psi\ w,$$

where $\varphi = \angle(\delta_{i+1}, z_i) \in [0, \pi]$ is the polar angle with respect to $e_1'$, $\psi \in [0, 2\pi)$ is the azimuthal angle around $e_1'$ with respect to $u$, and $w$ is a unit vector in $(e_1', u)^\perp$.

##### 4.2 Projection of $\delta_i$ onto the local frame

We compute the projections of $\delta_i$ onto $e_1'$ and $u$.

For the first projection, since $\langle \delta_i,\, e_1'\rangle = \langle \delta_i,\, z_i\rangle/d_i$, and $z_i = z_{i-1} + \delta_i$ with $\langle z_{i-1},\, \delta_i\rangle = r_i\, d_{i-1}\cos\theta_i$:

$$\langle \delta_i,\, z_i\rangle = \langle \delta_i,\, z_{i-1}\rangle + \|\delta_i\|^2 = r_i\, d_{i-1}\cos\theta_i + r_i^2.$$

For the second projection, using $\|u\| = 1$, $u \perp e_1'$, and the explicit expression of $u$:

$$\langle \delta_i,\, u\rangle = \frac{1}{d_i}\big(-r_i^2\sin\theta_i\cdot\cos\theta_i + (d_{i-1} + r_i\cos\theta_i)\cdot r_i\sin\theta_i\big) = \frac{d_{i-1}\, r_i\sin\theta_i}{d_i}.$$

These two results give:

$$\langle \delta_i,\, e_1'\rangle = \frac{r_i^2 + r_i\, d_{i-1}\cos\theta_i}{d_i}, \qquad \langle \delta_i,\, u\rangle = \frac{d_{i-1}\, r_i\sin\theta_i}{d_i}. \tag{29}$$

##### 4.3 Closed formula for epistasis between consecutive beneficial mutations

Substituting (29) into the expression of $\langle \delta_i,\, \delta_{i+1}\rangle$ :
Expanding the inner product using the frame $(e_1', u, w)$:

$$\langle \delta_i,\, \delta_{i+1}\rangle = \langle \delta_i,\, e_1'\rangle\cdot r_{i+1}\cos\varphi + \langle \delta_i,\, u\rangle\cdot r_{i+1}\sin\varphi\cos\psi + \underbrace{\langle \delta_i,\, w\rangle}_{=0}\cdot r_{i+1}\sin\varphi\sin\psi,$$

where the last term vanishes since $w \in (e_1', u)^\perp$ and $\delta_i$ lies in the plane $(e_1', u)$ by construction.We get :

$$\langle \delta_i,\, \delta_{i+1}\rangle = \frac{r_i^2 + r_i\, d_{i-1}\cos\theta_i}{d_i}\cdot r_{i+1}\cos\varphi + \frac{d_{i-1}\, r_i \sin\theta_i}{d_i}\cdot r_{i+1}\sin\varphi\cos\psi. \tag{30}$$

Since $\delta_{i+1} \sim \mathcal{N}(0, \sigma^2 I_n)$ is isotropic, $\psi$ is independent of $(r_{i+1}, \varphi)$ and uniform on $[0, 2\pi)$, hence $\mathbb{E}[\cos\psi] = 0$. Taking expectations:

$$\mathbb{E}\big[\langle \delta_i,\, \delta_{i+1}\rangle\big] = \mathbb{E}\left[\underbrace{\frac{r_i^2 + r_i\, d_{i-1}\cos\theta_i}{d_i}}_{=:\, P_i}\cdot \underbrace{\mathbb{E}_{f_{\mathrm{B}}(\cdot|d_i)}\big[r_{i+1}\cos\varphi\big]}_{=:\, Q_B(d_i)}\right]. \tag{31}$$

$P_i$ is the projection of $\delta_i$ onto the radial direction $e_1' = z_i/d_i$ after fixation, and $Q_B(d_i)$ is the mean radial projection of the next beneficial mutation from distance $d_i$. The key observation is that $(r_{i+1}, \varphi)$, the norm and polar angle of $\delta_{i+1}$, depend on $(r_i, \theta_i)$ only through $d_i = d_i(r_i, \theta_i)$: the next mutation sees only the current distance to the optimum, not the details of how it was reached

#### 4.4 Regime I: asymptotic computation

##### 4.4.1 New distance to the optimum after fixation of $\delta_i$

For fixed $(r_i, \theta_i)$, the new distance to the optimum is $d_i := d(r_i, \theta_i) = \sqrt{d_{i-1}^2 + 2r_i d_{i-1}\cos\theta_i + r_i^2}$. In Regime I, the ratio $r_i/d_{i-1} \sim \tilde{\sigma}/d_{i-1} = 2x/\sqrt{n}$ is small for large $n$. Using concentration with $n$, $r_i \sim \tilde{\sigma}$ and $\sqrt{n}\cos\theta_i = O(1)$ :

$$\frac{d_i}{d_{i-1}} = \sqrt{1 + \underbrace{\frac{2r_i\cos\theta_i}{d_{i-1}}}_{O(1/n)} + \underbrace{\frac{r_i^2}{d_{i-1}^2}}_{O(1/n)}} = 1 + O\left(\frac{1}{n}\right).$$

Therefore the Fisher parameter at the next step satisfies:

$$\frac{n\sigma}{2d_i} = x\cdot\frac{d_{i-1}}{d_i} \underset{n\to\infty}{\longrightarrow} x.$$

In high dimension, each mutation is negligibly small relative to the distance to the optimum: the next beneficial mutation sees the same Fisher parameter $x$, independently of the details of the previous step.

##### 4.4.2 Computation of $Q_B(d_i)$ and main result in regime I

By exactly the same calculation as for Term $B$ in section 3.2, applied from distance $d_i$:

$$Q_B(d_i) = \mathbb{E}_{f_{\mathrm{B}}(\cdot|d_i)}[r_{i+1}\cos\varphi] \underset{n\to\infty}{\sim} -\sigma\,\Lambda\left(\frac{n\sigma}{2d_i}\right).$$

Since $n\sigma/(2d_i) \to x$ by the previous section, we obtain:

$$Q_B(d_i) \underset{n\to\infty}{\sim} -\sigma\,\Lambda(x). \tag{32}$$

By exactly the same calculation as for Term $B$ in section 3.2, applied from distance $d_i$, the Laplace method gives concentration at $t^* = 1$, and using (5) and (6):

$$Q_B(d_i) \underset{n\to\infty}{\sim} -\frac{\tilde{\sigma}\, d_i}{\sqrt{n}} \cdot \frac{e^{-x^2/2}}{\displaystyle\int_x^\infty e^{-v^2/2}\, dv} = -\sigma\, \Lambda(x).$$

Again with the Laplace method, the full expectation assembles as:

$$\mathbb{E}\big[\langle \delta_i,\, \delta_{i+1}\rangle\big] \underset{n\to\infty}{\sim} \underbrace{\frac{\sqrt{n}}{\displaystyle\int_x^\infty e^{-v^2/2}\, dv}}_{\text{from } K_B} \cdot \underbrace{\frac{-2x}{n}\,\frac{e^{-x^2/2}}{\displaystyle\int_x^\infty e^{-v^2/2}\, dv}}_{\text{from } Q_B/d(r,\theta)} \cdot \underbrace{\left[\frac{\tilde{\sigma}^2}{\sqrt{n}}\int_x^\infty e^{-\frac{v^2}{2}}\, dv - \frac{\tilde{\sigma}\sqrt{n}\,\tilde{\sigma}}{2x} \cdot \frac{1}{n}\, e^{-x^2/2}\right]}_{\text{from } P_i}.$$

$$\mathbb{E}\big[\langle \delta_i,\, \delta_{i+1}\rangle\big] \underset{n\to\infty}{\sim} \sigma^2\, \Lambda(x)\big(\Lambda(x) - 2x\big) = \frac{4d^2x^2}{n^2}\Lambda(x)\big(\Lambda(x) - 2x\big). \tag{33}$$

The sign of (33) is determined by $\Lambda(x) - 2x$: this changes sign at the unique $x^* \approx 0.612$, solution of $\Lambda(x^*) = 2x^*$. Consecutive mutations are positively correlated for $x < x^*$ and negatively correlated for $x > x^*$.

#### 4.5 Regime II: asymptotic computation

In Regime II, $y := \tilde{\sigma}/(2d_{i-1}) = O(1)$ and $x = y\sqrt{n} \to \infty$. Note that $y < 1$ since beneficial mutations require $\tilde{\sigma} < 2d_{i-1}$.

##### 4.5.1 Computation of $Q_B$

For a given $d = d(r, \theta)$, we introduce $q := \tilde{\sigma}/(2d) = O(1)$, which plays the role of $y$ but at distance $d$ rather than $d_{i-1}$. Setting $r = \tilde{\sigma}\, t$, we write $Q_B(d) = N/D$ with:

$$N = -\tilde{\sigma} \int_0^{1/q} t^n\, e^{-nt^2/2} \int_{\arccos(-qt)}^{\pi} \sin^{n-2}\varphi \cos\varphi\, d\varphi\, dt,$$
$$D = \int_0^{1/q} t^{n-1}\, e^{-nt^2/2} \int_{\arccos(-qt)}^{\pi} \sin^{n-2}\varphi\, d\varphi\, dt. \tag{34}$$

Setting $F(\varphi) := \log\sin\varphi$, the Laplace method at the edge gives, for any fixed $\gamma \in (\pi/2, \pi)$:

$$\int_\gamma^\pi e^{(n-2)F(\varphi)}\, d\varphi \underset{n\to\infty}{\sim} \frac{(\sin\gamma)^{n-2}}{(n-2)|\cos\gamma|},$$

since $F'(\gamma) = \cos\gamma/\sin\gamma < 0$. For the integral with $\cos\varphi$, the exact primitive gives directly:

$$\int_\gamma^\pi \sin^{n-2}\varphi \cos\varphi\, d\varphi = -\frac{(\sin\gamma)^{n-1}}{n-1}.$$

Substituting $\gamma = \arccos(-qt)$, so that $\sin\gamma = \sqrt{1 - q^2t^2}$ and $|\cos\gamma| = qt$, the angular factors combine and the ratio $N/D$ simplifies to:

$$Q_B \underset{n\to\infty}{\sim} -\tilde{\sigma} q \frac{\displaystyle\int_0^{1/q} t^n\, e^{-nt^2/2}\big(1 - q^2t^2\big)^{(n-1)/2}\, dt}{\displaystyle\int_0^{1/q} t^{n-2}\, e^{-nt^2/2}\big(1 - q^2t^2\big)^{(n-1)/2}\, dt}.$$

Factoring $t^2$ from the numerator, both integrals share the same integrand, so the ratio equals $\mathbb{E}[t^2]$ under the Laplace measure, which concentrates at $t^*$:

$$Q_B \sim -\tilde{\sigma} q \cdot (t^*)^2 = -2dq^2\,(t^*)^2,$$

where:

$$t^* = \underset{t\in(0,1/q)}{\operatorname{argmax}} \left[\log t - \frac{t^2}{2} + \frac{1}{2}\log(1-q^2t^2)\right].$$

Differentiating and setting to zero:

$$y^2t^4 - (1+2q^2)t^2 + 1 = 0,$$

with root in $(0, 1/q)$:

$$(t^*)^2 = \frac{(1+2q^2) - \sqrt{1+4q^4}}{2q^2}.$$

$$\frac{Q_B(d)}{d} \underset{n\to\infty}{\sim} -\left(1+2q^2-\sqrt{1+4q^4}\right), \qquad q := \frac{\tilde{\sigma}}{2d}. \tag{35}$$

###### 4.5.2 Main result in regime II

Recall that $q := \tilde{\sigma}/(2d)$ where $d = d(r,\theta)$. Since $d(r,\theta) = d_{i-1}\sqrt{1 + 2r\cos\theta/d_{i-1} + r^2/d_{i-1}^2}$, we have:

$$q = \frac{\tilde{\sigma}}{2d(r,\theta)} = y\,\frac{d_{i-1}}{d(r,\theta)} = \frac{y}{\sqrt{1+4yt\cos\theta+4y^2t^2}},$$

where we used the change of variable $r = \tilde{\sigma}\,t$, giving $r/d_{i-1} = 2yt$, together with $y = \tilde{\sigma}/(2d_{i-1})$. The full expectation then writes:

$$\frac{1}{\tilde{\sigma}^2}\,\mathbb{E}\big[\langle\delta_i,\,\delta_{i+1}\rangle\big] = \frac{\displaystyle\int_0^{1/y} dt\,t^{n-1}\,e^{-\frac{nt^2}{2}} \int_{\arccos(-yt)}^{\pi} d\theta\,\sin^{n-2}\theta\left[t^2 + \frac{t\cos\theta}{2y}\right]\frac{Q_B}{d}(t,\theta)}{\displaystyle\int_0^{1/y} dt\,t^{n-1}\,e^{-\frac{nt^2}{2}} \int_{\arccos(-yt)}^{\pi} d\theta\,\sin^{n-2}\theta}, \tag{36}$$

For any $t = O(1)$ as $n\to\infty$ (ie not going to 0), the angular integral concentrates at the edge $\theta = \arccos(-yt)$ by the same Laplace argument as above, giving:

$$\int_{\arccos(-yt)}^{\pi} \sin^{n-2}\theta\,d\theta \underset{n\to\infty}{\sim} \frac{1}{nyt}\,(1-y^2t^2)^{\frac{n-1}{2}}.$$

At the edge $\cos\theta = -yt$, we have:

$$\left[t^2 + \frac{t\cos\theta}{2y}\right]_{\cos\theta=-yt} = t^2 - \frac{t^2}{2} = \frac{t^2}{2}.$$

So we get:

$$\frac{1}{\tilde{\sigma}^2}\,\mathbb{E}\big[\langle\delta_i,\,\delta_{i+1}\rangle\big] \underset{n\to\infty}{\sim} \frac{\displaystyle\int_0^{1/y} t^{n-1}\,e^{-nt^2/2}\cdot(1-y^2t^2)^{(n-1)/2}\cdot 1/(nyt)\cdot\frac{t^2}{2}\,dt}{\displaystyle\int_0^{1/y} t^{n-1}\,e^{-nt^2/2}\cdot(1-y^2t^2)^{(n-1)/2}\cdot 1/(nyt)\,dt} \times \frac{Q_B}{d}$$

Since $\cos\theta$ concentrates near $-yt$ at the edge, we have $q \sim y$, i.e. $\tilde{\sigma}/(2d(r,\theta)) \sim \tilde{\sigma}/(2d_{i-1})$: the distance to the optimum remains approximately constant. This has a clear biological interpretation: in Regime II, beneficial mutations are so rare that only those arising near the boundary of beneficiality $\cos\theta \approx -r/(2d_{i-1})$ contribute, and these mutations leave the population on approximately the same sphere around the optimum.

Consequently, $Q_B/d$ is evaluated at $q = y$, giving by (35):

$$\frac{Q_B}{d} \underset{n\to\infty}{\sim} -\left(1 + 2y^2 - \sqrt{1+4y^4}\right). \tag{37}$$

By the Laplace method, $t$ concentrates around the same concentration point as previously.

$$t^* = \underset{t\in(0,1/q)}{\operatorname{argmax}} \left[\log t - \frac{t^2}{2} + \frac{1}{2}\log(1-q^2t^2)\right].$$

Finally:

$$\mathbb{E}\big[\langle \delta_i,\, \delta_{i+1}\rangle\big] \underset{n\to\infty}{\sim} -\frac{\tilde{\sigma}^2}{4y^2}\left(1 + 2y^2 - \sqrt{1+4y^4}\right)^2. \tag{38}$$

##### 4.6 Comparison of the two regimes and matching

**Sign of the correlation.** In Regime I ($x = O(1)$), the expectation $-\mathbb{E}[\langle \delta_i,\, \delta_{i+1}\rangle] = \sigma^2\Lambda(x)(2x - \Lambda(x))$ changes sign at $x^* \approx 0.612$: consecutive mutations have positive epistasis close to the optimum and negative epistasis far from it. In Regime II ($y = O(1)$), the average epistasis $\tilde{\sigma}^2(1 + 2y^2 - \sqrt{1+4y^4})^2/(4y^2)$ is *always positive*, reflecting the fact that beneficial mutations near the boundary of beneficiality always tend to require compensation.

**Scaling with $n$.** In Regime I with $x$ fixed, $\mathbb{E}[\langle \delta_i,\, \delta_{i+1}\rangle] = O(\tilde{\sigma}^2/n) \to 0$: the correlation vanishes as $n \to \infty$ under the combined effect of two reinforcing mechanisms. First, as $d \sim D\sqrt{n} \to \infty$, the beneficiality condition $\cos\theta < -\tilde{\sigma}/(2D\sqrt{n}) \to 0$ becomes increasingly permissive: the radial component required for a mutation to be beneficial becomes negligibly small, so beneficial mutations carry almost no directional information about the optimum. Second, as $n$ grows, the orthogonal complement of the radial axis has dimension $n-1 \to \infty$, and by spherical concentration (section 1.3), $\delta_{i+1}$ lies almost surely in this orthogonal subspace: consecutive mutations are asymptotically orthogonal (same argument), and their inner product vanishes.

In Regime II ($y = O(1)$, $d = O(1)$), we have $\mathbb{E}[\langle \delta_i,\, \delta_{i+1}\rangle] = O(\tilde{\sigma}^2) = O(1)$: the correlation remains of order one regardless of $n$. This reflects the fact that in this regime, the geometry is essentially $n$-independent — each mutation is of the same order as the distance to the optimum and geometric constrain is very strong so mutations lies in the boundary of beneficial ball.

**Matching of the two regimes.** The two asymptotic expressions are consistent in their common limit $x \to \infty$ (Regime I) and $y \to 0$ (Regime II), with $y = x/\sqrt{n}$.

In Regime I as $x \to \infty$, using $\Lambda(x) \sim x$:

$$\mathbb{E}\big[\langle \delta_i,\, \delta_{i+1}\rangle\big] \underset{x\to\infty}{\sim} -\sigma^2x^2 = -\frac{\tilde{\sigma}^2x^2}{n}.$$

In Regime II as $y \to 0$, using $1 + 2y^2 - \sqrt{1+4y^4} \sim 2y^2$:

$$\mathbb{E}\big[\langle \delta_i,\, \delta_{i+1}\rangle\big] \underset{y\to 0}{\sim} -\frac{\tilde{\sigma}^2}{4y^2}(2y^2)^2 = -\tilde{\sigma}^2y^2 = -\frac{\tilde{\sigma}^2x^2}{n}.$$

Both limits coincide, confirming the consistency of the two asymptotic expansions in the intermediate regime $y \to 0$, $x \to \infty$, $y\sqrt{n} = x = O(\sqrt{n})$.

**Properties of the Regime II correlation** The function $h(y) := \dfrac{1}{4y^2}\left(1 + 2y^2 - \sqrt{1 + 4y^4}\right)^2$ is always positive on $(0, 1)$, with:

- $h(y) \to 0$ as $y \to 0$;
- $h(1) = \frac{1}{4}(3 - \sqrt{5})^2 \approx 0.146$;
- a unique maximum at $y^* = 1/\sqrt{2}$, which can be verified analytically by setting $h'(y) = 0$ and checking that $y = 1/\sqrt{2}$ satisfies the stationarity condition exactly, with $h(1/\sqrt{2}) = \frac{1}{2}(2 - \sqrt{2})^2/2 = (3 - 2\sqrt{2})/2 \approx 0.172$.

Therefore $\mathbb{E}[\langle \delta_i,\, \delta_{i+1} \rangle]/\tilde{\sigma}^2 = -h(y)$ is always negative, reaches its most negative value $\approx -0.172\,\tilde{\sigma}^2$ at $y^* \approx 1/\sqrt{2}$, and satisfies $|\mathbb{E}[\langle \delta_i,\, \delta_{i+1} \rangle]| = O(\tilde{\sigma}^2) = O(1)$: the anti-correlation is strong and of order one throughout Regime II.

### 5 Supplementary figures

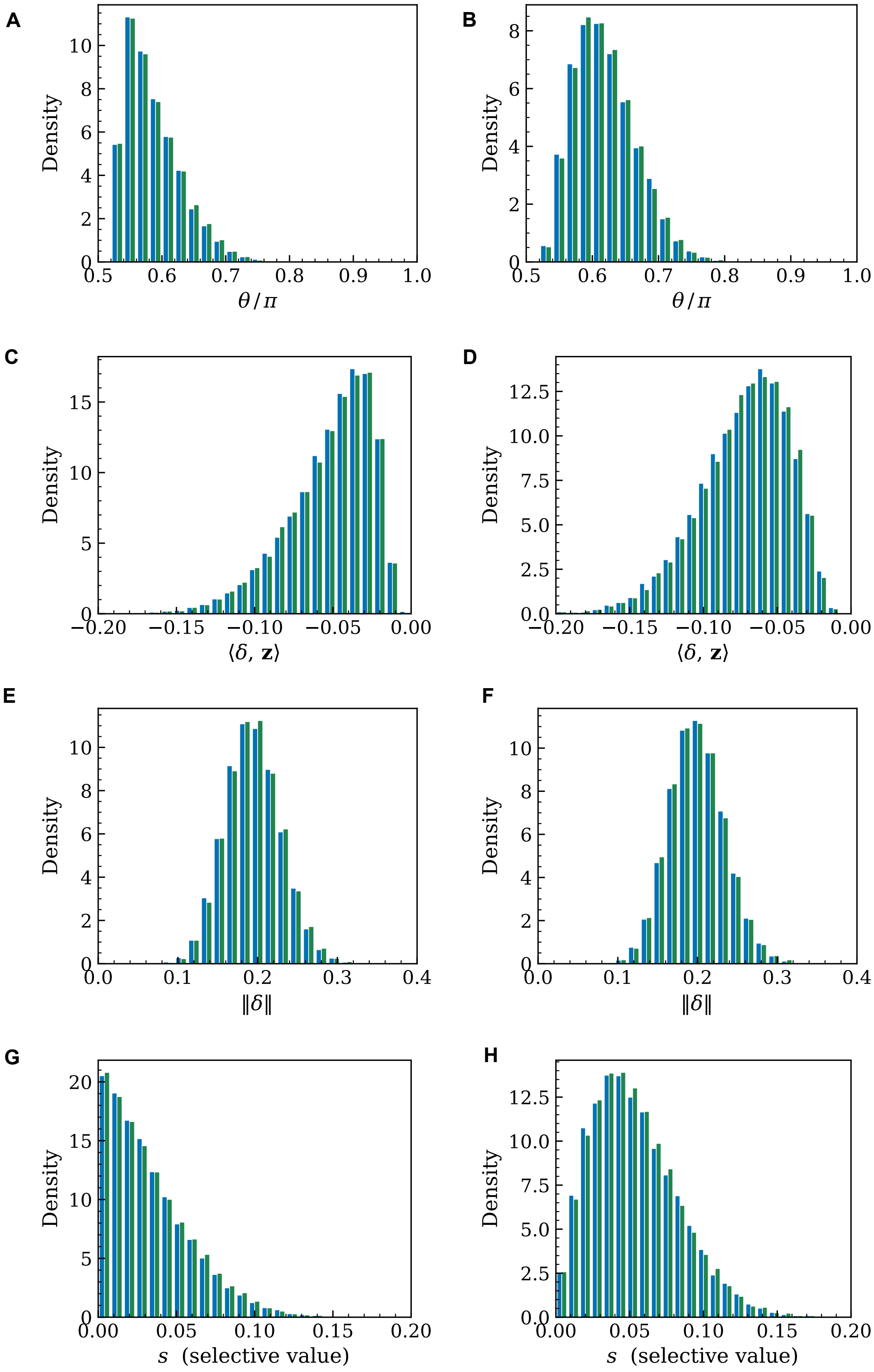
A
B
C
D
E
F
G
H
Density
θ / π
⟨δ, z⟩
‖δ‖
s (selective value)

Figure S1: **Validity of the Metropolis-Hastings Markov Chain Monte Carlo (MCMC) approach for sampling fixed mutations in Fisher's geometric model.** Each panel compares the distribution of mutational properties for 20,000 vectors obtained by MCMC-Metropolis-Hastings sampling (blue) and by naive rejection sampling (green), the latter serving as ground truth. Left column: mutations accepted conditional on beneficiality only. Right column: mutations accepted with probability proportional to the Kimura fixation probability $p(s) = (1 - e^{-2s})/(1 - e^{-4N_e s})$, where $s$ is the selection coefficient and $N_e$ the effective population size. For each sampling regime, four quantities are shown: **(a-b)** the angle between the mutation and the selection axis, $\theta/\pi$; **(c-d)** the projection of the mutational vector onto the selection axis, $\langle \boldsymbol{\delta}, \mathbf{z} \rangle$; **(e-f)** the Euclidean norm of the mutation, $\|\boldsymbol{\delta}\|$; and**(g-h)** the selective value $s$. The near-perfect superposition of the two distributions across all panels and both regimes confirms that the MCMC-MH sampler correctly targets the conditional distribution of fixed mutations, including in parameter regimes where rejection sampling becomes computationally prohibitive. Simulations were performed with $n = 16$ phenotypic dimensions, initial distance to the optimum $d = 1$, mutational standard deviation $\sigma = 0.05$, and effective population size $N_e = 10^7$.

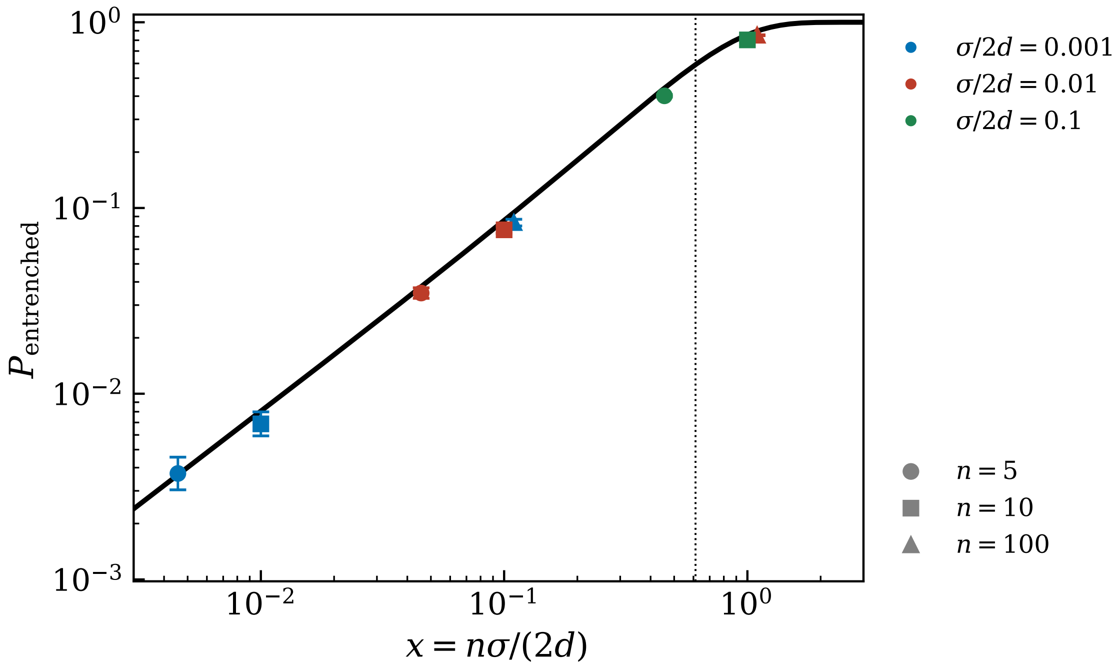

Figure S2: **The probability of entrenchment depends on the parameters $n$, $\sigma$, and $d$ only through the composite dimensionless parameter $x = n\sigma/(2d)$.** The solid black curve shows the theoretical prediction $P_{\text{entrenched}}(x) = [\Phi(2x) - \Phi(x)]/[1 - \Phi(x)]$, derived analytically for a beneficial mutation arising at distance $d$ from the optimum in a Fisher geometric model with $n$ phenotypic dimensions and mutational standard deviation $\sigma$ per axis. The vertical dotted line marks the critical threshold $x^* \approx 0.612$, below which diminishing returns epistasis predominates on average. Symbols show estimates from $2.5 \times 10^4$ simulated beneficial mutations for three values of $\sigma/(2d)$ (colours) and three values of $n$ (shapes), spanning a wide range of parameter combinations. The near-perfect collapse of all simulation points onto the theoretical curve confirms that $P_{\text{entrenched}}$ is a function of $x$ alone, independent of $n$, $\sigma$, and $d$ individually. Error bars show 95% Wilson score confidence intervals.

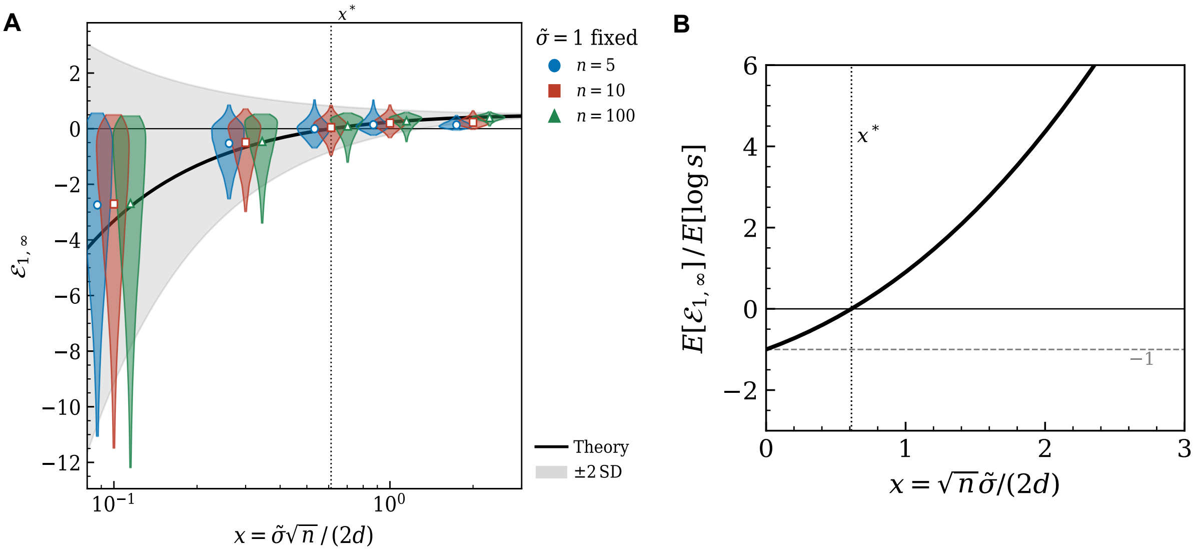

Figure S3: **Expected entrenchment of the first substitution once the adaptive walk has reached the optimum ($\mathcal{E}_{1,\infty}$) in Fisher's geometric model.** **(a)** Generalized epistasis $\mathcal{E}_{1,\infty}$ as a function of the dimensionless parameter $x = \tilde{\sigma}\sqrt{n}/(2d)$ (log scale), with $\tilde{\sigma} = 1$ fixed and $d$ varying to span the range of $x$ values. The solid black curve shows the asymptotic expectation $\mathbb{E}[\mathcal{E}_{1,\infty}] \sim \tilde{\sigma}^2(1 - \Lambda(x)/2x)$ and the shaded band its $\pm 2$ standard deviations, where $\Lambda(x) = \phi(x)/[1 - \Phi(x)]$ is the inverse Mills ratio. At fixed $\tilde{\sigma}$, both quantities depend on $n$, $\sigma$, and $d$ only through $x$, so that the theoretical prediction is universal. The vertical dotted line marks $x^* \approx 0.612$, defined by $\Lambda(x^*) = 2x^*$, at which $\mathbb{E}[\mathcal{E}_{1,\infty}]$ changes sign: below $x^*$ the first substitution experiences diminishing returns on average, while above $x^*$ it is entrenched on average. Violin plots show the empirical distribution of $\mathcal{E}_{1,\infty}$ from 25,000 simulated beneficial mutations for each combination of $d$ ($x$-axis position) and $n$ (colours, jittered horizontally for clarity). The collapse of all violins onto the single theoretical curve confirms that, at fixed $\tilde{\sigma}$, $\mathcal{E}_{1,\infty}$ depends on the biological parameters only through $x$. **(b)** Ratio of the expected generalized epistasis to the expected log-fitness gain of the same substitution, $\mathbb{E}[\mathcal{E}_{1,\infty}]/\mathbb{E}[\log s]$, as a function of $x$. Using $\mathbb{E}[\log s] \sim (2d^2x/n)(\Lambda(x) - x)$, the $n$- and $\sigma$-dependent prefactors cancel exactly, yielding the universal ratio $(2x - \Lambda(x))/(\Lambda(x) - x)$. This ratio is always negative for $x < x^*$, equals $-1$ at $x = 0$ (dashed gray line), and diverges as $x^2$ for large $x$. It provides a direct measure of the relative magnitude of path-dependent epistasis: a ratio of $-1$ indicates that the epistatic cost of reversion at the optimum exactly equals the original selective benefit, while large positive values indicate that the substitution has become deeply entrenched relative to its initial benefit.

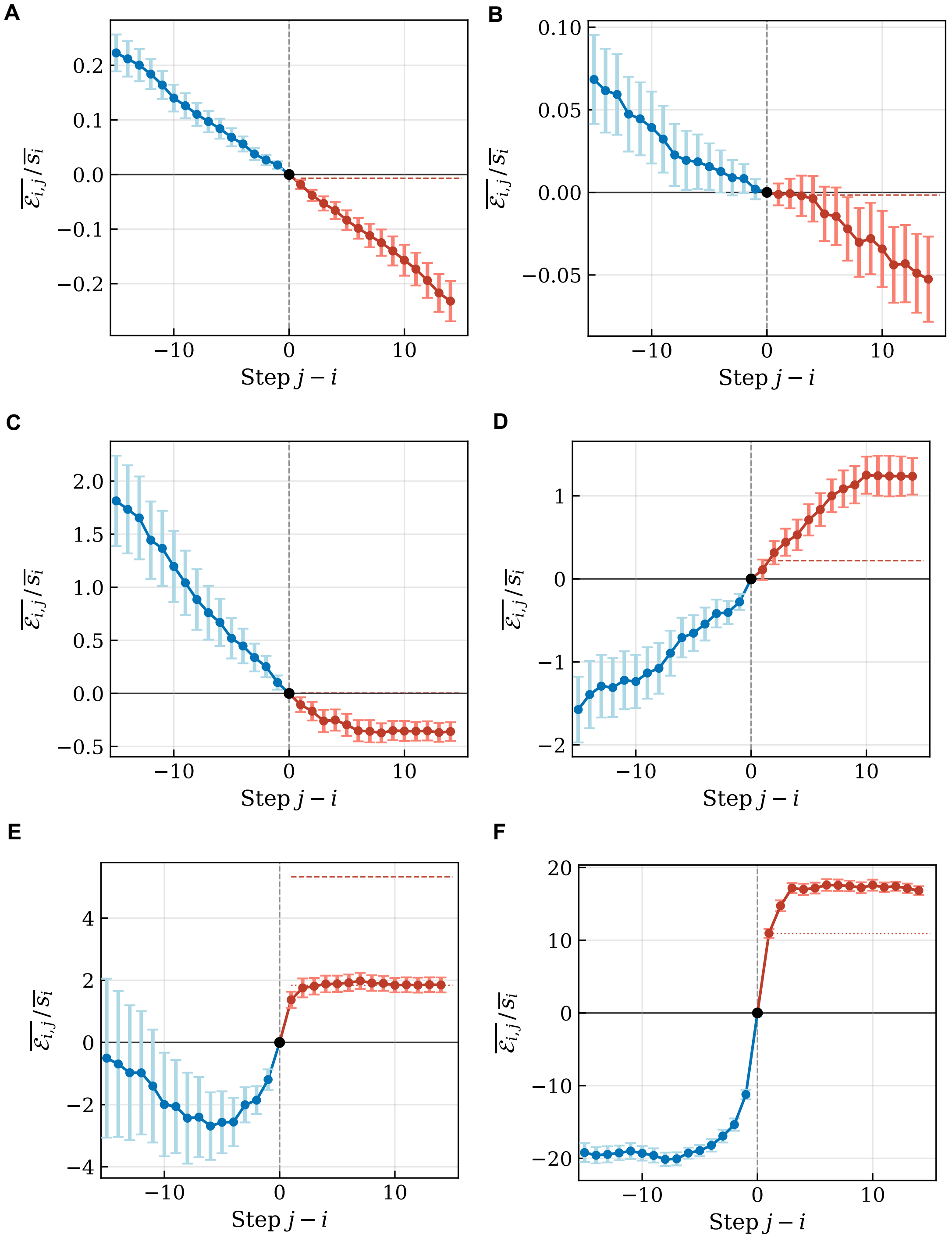
A
B
C
D
E
F
$\overline{\varepsilon_{i,j}}/\overline{s_i}$
Step $j-i$

Figure S4: **Contingency and entrenchment of a focal substitution along adaptive walks in Fisher's geometric model, with fixation probability weighting.** This Figure is the same as Figure 3, except that each mutation is accepted with probability proportional to the Kimura fixation probability (equation 8), rather than uniformly among beneficial mutations. Columns correspond to phenotypic complexity ($n = 10$, panels **a**–**c**–**e**; $n = 100$, panels **b**–**d**–**f**); rows correspond to decreasing distance to the optimum ($d = 2.5$, panels **a**,**b**; $d = 0.25$, panels **c**,**d**; $d = 0.07$, panels **e**,**f**), with $\tilde{\sigma} = 0.1$ fixed throughout. The focal substitution is the 16th step of each walk, standardized to occur at distance $d$ from the optimum (relative tolerance 10%). For each panel, 200 independent adaptive walks were simulated under the SSWM regime with Kimura fixation probability weighting. Blue circles show the mean generalized epistasis coefficient $\overline{\mathcal{E}_{i,j}}/\overline{s_i}$ for $j < i$ (contingency) and red circles for $j > i$ (entrenchment), normalized by the mean log-fitness gain $\overline{s_i}$ of the focal substitution. Error bars show $\pm 2$ standard errors across replicates. The black dot at $j - i = 0$ marks the focal substitution ($\mathcal{E}_{i,i} = 0$ by definition). The dashed line shows the Regime I asymptotic prediction for $\mathbb{E}[\mathcal{E}_{i,i+1}]/\overline{s_i}$ (equation 33); the dotted line shows the Regime II prediction (equation 38, displayed only when $y \geq 0.3$). Dimensionless parameters for each panel: **(a)** $y = 0.02$, $x = 0.063$; **(b)** $y = 0.02$, $x = 0.2$; **(c)** $y = 0.20$, $x = 0.63$; **(d)** $y = 0.20$, $x = 2$; **(e)** $y = 0.71$, $x = 2.26$; **(f)** $y = 0.71$, $x = 7.1$. The critical threshold $x^* \approx 0.612$ is predicted to separate diminishing returns ($x < x^*$) from entrenchment ($x > x^*$). Notably, panel **(c)** shows that, near the transition $x \approx x^*$, weighting by fixation probability preserves a diminishing returns regime, in contrast to the unweighted case in Figure 3. This reflects the fact that weighting by fixation probability favors larger mutations more strongly aligned with the direction of the optimum (see Figure S1), shifting the effective distribution of fixed mutations toward better-aligned vectors and thereby delaying the transition to entrenchment to a slightly higher threshold on $x$.
